# Accessibility-guided targeting of NRF2-responsive long noncoding RNA NMRAL2P alters a distinct gene expression program and reduces non-small cell lung cancer proliferation

**DOI:** 10.64898/2026.11.13.765723

**Authors:** Jae A. Bucknor, Brittany M. Bowman, Shannon L. Stein, Chase A. Weidmann

**Author notes:** Jae A. Bucknor and Brittany M. Bowman contributed equally to this work.

## Abstract

Aberrant activation of the antioxidant transcription factor NRF2 promotes non-small cell lung cancer (NSCLC) survival and therapy resistance, but downstream effectors beyond redox regulation remain incompletely defined. Transcriptome profiling of lung and NSCLC cells identified the pseudogene-derived long noncoding RNA LOC344887/NMRAL2P as consistently associated with NRF2 activity. The *NMRAL2P* locus produces three predominant, nuclear-localized isoforms whose expression depends on NRF2 across normal lung, lung adenocarcinoma, and lung squamous carcinoma cells. In-cell SHAPE-MaP identified accessible regions within NMRAL2P transcripts, guiding antisense gapmer design. NMRAL2P depletion inhibited NSCLC cell proliferation and increased sensitivity to cisplatin. Transcriptome analysis revealed that NMRAL2P depletion altered a restricted gene expression program involving cell adhesion and MAPK signaling, with responses distinct from those observed following NRF2 depletion. Core NRF2 antioxidant target genes were largely unaffected by NMRAL2P depletion. These findings support a model in which NRF2 induces NMRAL2P to sustain a distinct gene expression program that contributes to NSCLC proliferation and survival. NMRAL2P represents an RNA effector downstream of NRF2 that links oncogenic NRF2 activity to cellular functions beyond canonical antioxidant responses, providing a potential point of intervention in NRF2-active lung cancers and motivating further investigation of the mechanisms connecting this nuclear RNA to signaling pathways.

Graphical Abstract

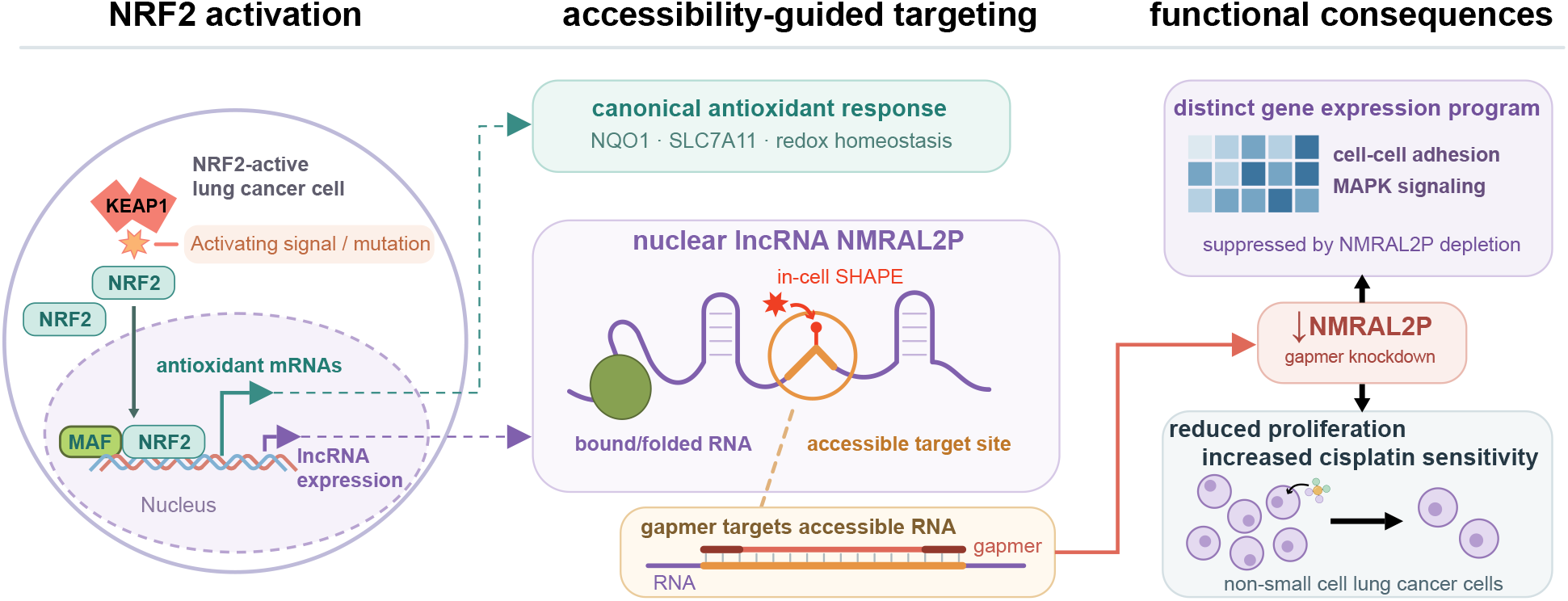

## INTRODUCTION

Lung cancer remains the leading cause of cancer mortality worldwide (**1**). Although lung cancer incidence has declined in many populations as smoking has decreased (**2, 3**), ambient air pollution and residential radon remain important causes of disease (**4–7**). In 2022, an estimated 195, 000 new cases of lung adenocarcinoma worldwide were attributable to ambient particulate matter, with the greatest burden in East Asia (**3**). As electronic cigarette use becomes more common in young adults (**8**), its long-term effects on lung cancer incidence remain uncertain (**9, 10**). In the United States, exposure to major risk factors is unevenly distributed: smoking is more prevalent in socioeconomically disadvantaged communities (**11**), and communities of color experience disproportionate exposure to ambient fine particulate matter (PM_2.5_) (**12, 13**). Residential radon levels vary with local geology and building characteristics, and testing for elevated indoor radon is less common in socioeconomically disadvantaged neighborhoods (**14**). The current toll of lung cancer and the persistence of its major risk factors underscore the need for more effective treatments. Non-small cell lung cancer (NSCLC), which includes lung adenocarcinoma and lung squamous cell carcinoma, accounts for approximately 85% of lung cancer diagnoses (**15**). A substantial fraction (∼25%) of NSCLC tumors exhibit aberrant activation of nuclear factor erythroid 2-related factor 2 (NRF2) (**16, 17**), a transcription factor that coordinates cellular responses to oxidative stress (**18**).

The KEAP1-NRF2 pathway couples the cellular response to oxidative and electrophilic stress to the rate of NRF2 degradation. Under basal conditions, Kelch-like ECH-associated protein 1 (KEAP1) forms a homodimer and serves as the substrate adaptor for a CUL3-RBX1 E3 ubiquitin ligase (**19, 20**). The BTB region of KEAP1 mediates dimerization and contributes to CUL3 recruitment, while the Kelch domains of the two KEAP1 subunits engage two motifs in the Neh2 domain of a single NRF2 molecule. The higher-affinity ETGE motif anchors NRF2 to the dimer, and engagement of the lower-affinity DLG motif positions nearby NRF2 lysine residues for ubiquitination (**21, 22**). The proteasome rapidly degrades ubiquitinated NRF2, keeping its basal abundance low. While there are other minor routes for NRF2 turnover (**23, 24**), KEAP1 provides the principal and stress-sensitive strategy for NRF2 proteostasis.

KEAP1 senses stress through multiple reactive cysteines, with different residues contributing to the response to different stimuli (**25, 26**). Modifications can inhibit KEAP1 and NRF2 interaction (**27**) or uncouple KEAP1-NRF2 interaction from NRF2 ubiquitination (**28, 29**). For example, modification of Cys151 by the synthetic triterpenoid CDDO can interfere with KEAP1-CUL3 association (**20, 30**). As KEAP1-dependent turnover slows, newly synthesized NRF2 accumulates, enters the nucleus, and partners with small MAF proteins to bind antioxidant response elements in DNA promoters and activate transcription (**28, 31, 32**).

In NSCLC, recurrent mutations can bypass this stress-sensitive switch. Activating *NFE2L2* mutations cluster in and around the DLG and ETGE motifs, where they can disrupt substrate recognition, while loss-of-function *KEAP1* mutations impair the machinery that normally turns NRF2 over (**33, 34**). These mechanisms provide the rationale for comparing distinct modes of NRF2 activation in this study. We use the electrophilic triterpenoid bardoxolone methyl (CDDO-Me) for pharmacological activation and model persistent genetic activation by disrupting either NRF2 recognition with the ETGE-motif substitution E79Q or KEAP1 function through loss of *KEAP1*.

Active NRF2 limits damage caused by oxidative stress – like lipid peroxidation and protein oxidation – by coordinating pathways that modify and remove oxidants, restore oxidized thiols, and detoxify reactive products (**18, 35**). Among the NRF2-responsive proteins measured for validation here (**Fig. 1**): SLC7A11 imports cystine as a source of cysteine for reduced glutathione synthesis (**36, 37**); thioredoxin reductase 1 (TXNRD1) uses NADPH to regenerate reduced thioredoxin, which reduces disulfides in target proteins and supplies electrons to peroxiredoxins for peroxide removal (**38, 39**); NQO1 catalyzes two-electron reduction of quinones, limiting their ability to generate reactive oxygen species through redox cycling (**39, 40**); and SQSTM1/p62 is a selective autophagy adaptor that can reinforce NRF2 activity through negative regulation of KEAP1 (**41, 42**). Together, NRF2-regulated pathways increase NADPH production, support the synthesis and regeneration of reduced glutathione, and chemically modify reactive compounds to inactivate them and/or mark them for removal. NRF2 promotes the efflux of detoxification products, including glutathione conjugates, by upregulation of specific membrane transporters (**39, 43–45**).

**Figure 1:**
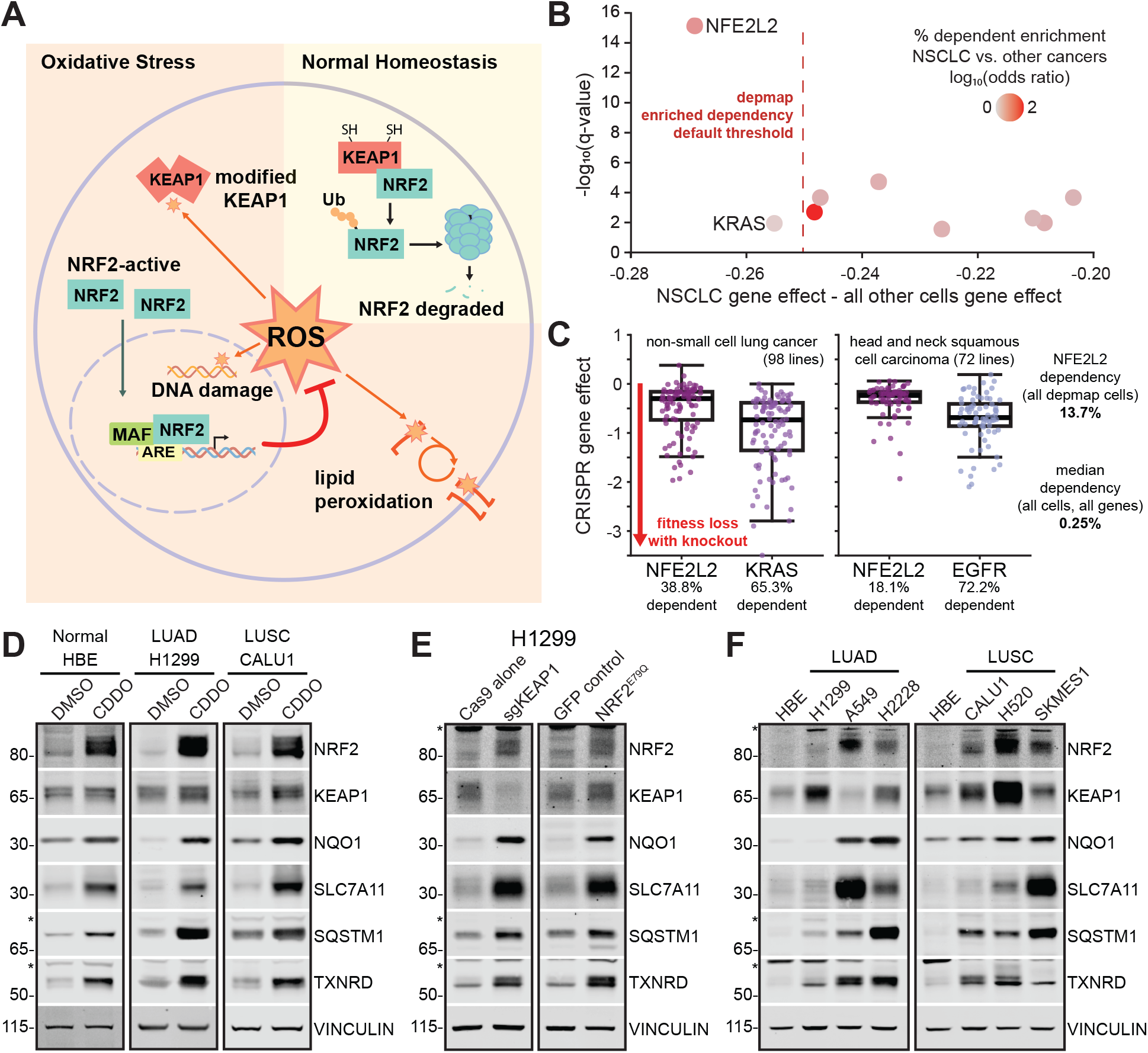
Disrupting KEAP1-NRF2 interaction activates NRF2 antioxidant signaling in lung and lung cancer cells. (**A**) Schematic of NRF2 proteostasis regulation by KEAP1 and NRF2 activation by oxidative stress (**B**) Cancer Dependency Map (26Q1) context explorer of non-small cell lung cancer cells comparing CRISPR deletion gene effect size and significance (q-value) in NSCLC cells and all other included cancer cell lines. The log odds ratio of the fraction of cell lines with dependency probabilities > 0.5 in each group (NSCLC versus other cancer lines) are represented by colors on each point. Default threshold = [Max. FDR ≤ 0.1, Min. |Effect Size| > 0.25, Min. frac of in-context lines dependent > 0.1]. (**C**) Gene effects for indicated cell types from knockout of NFE2L2 or cancer type-relevant gene (KRAS, EGFR). Boxes show the median and interquartile range; whiskers extend to the most extreme observations within 1.5× the interquartile range. Fraction of cell lines in each group with dependency probabilities > 0.5, the fraction of all DepMap cell lines dependent on NFE2L2, and the median dependency for all cells and all genes are displayed. Western blots showing NRF2, KEAP1, and NRF2 target protein levels comparing NRF2-inactive cells to (**D**) cells treated with CDDO-Me, (**E**) cells either expressing NRF2^E79Q^ or cells depleted of KEAP1 by CRISPR, and (**F**) cancer cells with elevated levels of NRF2 protein. Vinculin protein is a loading control. * label on western blots indicate non-specific bands.

While NRF2-activated antioxidant and detoxification programs have been extensively validated, sustained NRF2 activity can also influence other cell signaling behaviors relevant to tumor progression. In NSCLC cells, NRF2 has been linked to focal adhesion and motility through RhoA-ROCK signaling (**46**). Studies in other cellular contexts have also shown that NRF2 can suppress proinflammatory cytokine transcription and limit STING expression (**47, 48**). This broader activity raises the question of which features of the NRF2 response are shared across modes of activation and which downstream effectors could be targeted to inhibit lung cancer cell growth. Most studies have focused on protein-coding targets, although NRF2-responsive noncoding RNAs have also been identified, including NMRAL2P/LOC344887 (**49–51**).

NMRAL2P, also known as LOC344887, is a long noncoding RNA transcribed from a pseudogene derived from the protein-coding gene *NMRAL1*. Direct transcriptional activation of *NMRAL2P* by NRF2 has been demonstrated in colon cancer cells and lung cell models used to study pulmonary fibrosis (**50, 51**). Reported downstream functions include supporting NQO1 induction in sulforaphane-treated colon cancer cells and suppressing profibrotic gene expression in lung cells. In head and neck cancer models, NMRAL2P has also been implicated in promoting glycolysis and GPX2 transcription (**52**). These findings suggest that NMRAL2P can influence selected outputs of the NRF2 antioxidant response, with effects that depend on the cellular context. Higher NMRAL2P expression has been associated with poorer overall survival in NSCLC (**53**), but its contribution to NSCLC cell growth and its relationship to canonical NRF2-dependent antioxidant programs are not fully understood. Determining whether NMRAL2P supports established defenses or a distinct downstream programs could reveal additional mechanisms through which NRF2-active lung cancer cells sustain growth.

Here, we sought to identify shared features of NRF2-dependent gene expression and determine whether noncoding effectors contribute to the growth of NRF2-active NSCLC cells. We compared transcriptional responses across pharmacological and genetic models of NRF2 activation and NSCLC cells with constitutive pathway activity, identifying NMRAL2P among the consistently induced transcripts. We then characterized NMRAL2P transcript architecture and localization and used in-cell RNA structure probing to guide the design of antisense oligonucleotides for its depletion. By measuring proliferation and cisplatin sensitivity and comparing the transcriptional consequences of NMRAL2P and NRF2 depletion, we assessed whether NMRAL2P supports canonical antioxidant defenses or a distinct downstream response. Our findings support a model in which NMRAL2P sustains a gene-expression program associated with cell adhesion and growth signaling downstream of NRF2.

## MATERIAL AND METHODS

### DepMap cancer dependency analysis

DepMap Public 26Q1 data were downloaded (**54**), including context-explorer-NSCLC.csv (NSCLC-versus-out-group comparison export), CRISPRGeneEffect.csv (Chronos gene-effect scores), CRISPRGeneDependency.csv (dependency probabilities), and Model.csv (cell-line and cancer-type metadata). Model IDs were matched across files, and cancer-context membership was assigned using OncotreePrimaryDisease in Model.csv.

Outgroup comparisons were plotted directly from exported data. For each cancer context, individual cell-line Chronos scores for *NFE2L2* were plotted against those for the context-specific comparator gene. Boxes summarize the scores; points represent individual models. For the pan-cancer comparison, a model was classified as dependent when its DepMap dependency probability was > 0.5.

### Cell culture

Immortalized healthy lung cell line 16HBE14o-cells were purchased from Sigma while cancer cell lines A549, H1299, H2228, CALU1, SKMES1, H520 and immortalized kidney epithelial cells, HEK293T, were purchased from ATCC (**Supplementary Table S1**). In our Mutation cell category, modified H1299-eYFP-NQO1 (HYN) cells were used in place of parental H1299: a gift from M. Ben Major (**55**), these cells constitutively express nuclear-localized mCherry and harbor an eYFP cassette stably integrated into the first intron of the *NQO1* gene (**56**). Beyond expression of mCherry and eYFP, H1299 and HYN cells show no significant differences in expression, morphology, or NQO1 response. For cell line media requirements and reagents see **Supplementary Tables S2 and S3**. All cells were grown at 37 °C with 5% CO_2_. *KEAP1* CRISPR-based knockout H1299 cells (sgKEAP1) and mutant NRF2^E79Q^-expressing cells (H1299-NRF2^E79Q^) were generated by lentiviral infection of HYN cells and were selected using 10 µg/ml blasticidin S HCl and 1 µg/ml puromycin, respectively. Cells were passed a maximum of 10-times during experimentation and tested for mycoplasma routinely (Southern Biotech PCR kit). Unless specified as antibiotic-free or when including additional selection antibiotics, cells were grown in “general antibiotics” (10 units/mL penicillin and 10 μg/mL streptomycin).

### Construction of lentivirus plasmids

Lentiviral cargo vectors to generate NRF2-active H1299 cell lines (**Supplementary Table S4**) were either a gift from M. Ben Major (pHAGE-FLAG-NRF2-E79Q and pHAGE-UBC-Flag-GFP) or generated by Gibson assembly into parental plasmid pDECKO-mCherry (Addgene #78534). First, pDECKO-Cas9-Blast (**Supplementary File 1**) was generated by incorporating DNA encoding flag-tagged spCas9 protein and swapping a puromycin resistance gene for DNA encoding Blasticidin S Deaminase (BSD) protein. Cas9 and BSD are driven by the same EF1α promoter and expressed as a polycistronic transcript separated by a P2A site and whose transcription terminates with a woodchuck hepatitis virus posttranscriptional regulatory element (WPRE). From pDECKO-Cas9-Blast, digestion and Gibson assembly were used to create pSECKO-KEAP1-A-Cas9-Blast (**Supplementary File 2**) which added a *KEAP1*-targeting single-guide CRISPR RNA (sgRNA, spacer 5′ GCAGCCAGATCCCAGGCCTAG) under the control of a U6 promoter. Sequencing-verified annotated sequences for pDECKO-Cas9-Blast and pSECKO-KEAP1-A-Cas9-Blast are provided as supplementary material.

### Lentivirus production in HEK293T cells

Approximately 5 million HEK293T cells were plated in a 10 cm dish in growth media including general antibiotics and incubated at 37 °C. After 24 hours, media was aspirated and replaced with 10 mL of antibiotic-free growth media. For the transfection, a three-component viral plasmid system was transfected into the cells with FuGENE (Promega). Briefly, into plain DMEM media, 2 µg of the packaging plasmid (psPAX2, Addgene Plasmid #12260), 6 µg of the envelope (pMD2.G, Addgene Plasmid #12259), and 8 µg of lentiviral cargo plasmids. FuGENE (64 µl) was added directly to DNA mixtures and incubated for 10 minutes at room temperature. The entire transfection mixture was added dropwise to the 10 cm dish containing cells. After 24 hr, the media was removed and replaced with antibiotic-free DMEM containing 20% FBS, and cells were incubated for another 48 hr. Virus-containing media was extracted with a 20 mL syringe and passed through a 0.45 µm filter into spin concentration columns with a nominal molecular weight limit of 100 kDa. Virus was concentrated 10-fold (10 mL to 1 mL) before storage at −80 °C (long-term) or 4 °C (for immediate infections).

### Generation of stable NRF2-active cell lines using lentivirus

For each infection, 600, 000 HYN cells were plated in a 6 cm dish with 4 mL of antibiotic-free media (See **Supplementary Table S5** for all cell plating densities). Cells were incubated for 24 hr. Media was replaced with transduction media (antibiotic-free media and 10 µg/mL polybrene) and 100-150 µl of concentrated virus was added dropwise. After 24 hr, media was replaced with complete media to remove the virus. After 48 to 72 hours, media was replaced with antibiotic selection media and cells were monitored and split as necessary until expansion.

### Monoclonal isolation of *KEAP1* knockout and NRF2-E79Q cells

Polyclonal cell lines were then subjected to single cell plating by plating ∼100 cells per 15 cm dish and picking colonies using small 0.25% trypsin coated Whatman cloning disks (MilliporeSigma) to cover individual colonies. Tweezers were used to place the filter paper into 24-well dishes, and the filter paper was removed 24 hr later. Single cell colonies were expanded and tested by RT-qPCR (reverse transcription and quantitative polymerase chain reaction), western blotting, and genomic amplicon sequencing for final clone selection.

### Treatment with CDDO-Me

Cells were plated 24 hr before being treated with 100 nM of CDDO-Me or 0.1% DMSO. Cells were harvested 24 hr after treatment for protein or RNA as described below. For each cell line, see **Supplementary Table S5** for cell numbers used for plating.

### Protein extraction and visualization by western blotting

Cell lines were plated 24 hr before harvesting (see CDDO-Me treatment above for additional processing details) in 6-well plates. Cells were washed twice at room temperature in 1× PBS and lysed directly in-well using RIPA buffer (Invitrogen) supplemented with 1× Halt Protease and 1× Halt Phosphatase Inhibitor Cocktails (Thermo Fisher Scientific) and 250 units/mL of benzonase (Millipore Sigma). Lysed cells were moved to 1.5 ml tubes and incubated on ice for at least 40 minutes with occasional 5 second vortexing. Lysates were cleared by centrifugation at 4 °C for 15 minutes and stored at −80 °C.

Protein concentrations were quantified by BCA assay (Pierce). Protein lysates (10 µg) were prepared in 1× LDS buffer (Invitrogen) with 100 mM DTT and boiled at 70 °C for 7 minutes. Proteins were separated on a 4-12% Bis-Tris gel (NuPAGE Invitrogen) in 1× MOPS (Boston Bioproducts) along with a molecular weight ladder prepared in tandem (PageRuler™ Plus Prestained Protein Ladder (Thermo Fisher Scientific)). Proteins were transferred to 0.45 µM nitrocellulose membranes (Thermo Fisher Scientific) at 24 V for 1 hour in a 2× transfer buffer (NuPAGE™ Transfer Buffer (Invitrogen)) supplemented with 10% methanol. Membranes were blocked in 5% non-fat milk in 1× TBST [Tris-Buffered Saline (TBS, VWR) with 1% Tween-20] for 30 minutes at room temperature. After removing blocking buffer, primary antibody dilutions (in 1× TBST) were added to membranes and incubated overnight at 4 °C. After washing three times with 1× TBST, LI-COR fluorescent secondary antibody mixes (diluted in 5% non-fat milk in 1× TBST) were added to membranes and incubated at room temperature for 1 to 2 hours. Membranes were washed three times with 1× TBST and once with 1× TBS (no Tween-20). Blots were imaged with a LI-COR Odyssey M instrument (LI-COR). See **Supplementary Table S6** for western reagents and **Supplementary Table S7** for antibody information.

### RNA extraction and DNase treatment

85% confluent plates were washed twice in 1× PBS and RNA was recovered with 1 mL of TRIzol Reagent (Invitrogen) per 6-well or 10 cm dish (Corning) according to manufacturer’s protocol. Briefly, the upper, aqueous layer of a TRIzol/chloroform phase separation was precipitated with an equal volume of isopropanol, and the pellet was washed once with 75% ethanol before drying. Final RNA pellets were resuspended in 44 µL of RNase-free water. To remove contaminating DNA, 5 µL of 10× TURBO DNase buffer and 1 µL (2 units) of TURBO DNase enzyme (Invitrogen) were added to RNA and incubated at 37 °C. After 30 minutes, another 1 µL of TURBO DNase enzyme was added before incubating again at 37 °C for 30 minutes. DNA-free RNA was purified with Mag-Bind® Total Pure NGS SPRI beads (Omega Biotek) using a 1.8× ratio of bead mixture to sample. After a 5-minute incubation at room temperature, beads were captured on a magnetic rack, washed three times with 70% ethanol, and air-dried for 7 minutes before RNA was eluted in 20-50 µL of nuclease-free water. RNA integrity number (RIN) scores were measured for RNA samples on a TapeStation 4150 instrument (Agilent), and only values above 8 were processed for RNA sequencing (majority of RIN scores > 9). See **Supplementary Table S8** for RNA extraction reagents.

### Ribosomal RNA depletion

Ribosomal RNA (rRNA) was depleted from RNA samples using RNase H enzyme and a custom set of 130 rRNA-complementary DNA oligonucleotides (**Supplementary Table S9**). In a total of 17 µL, 5 µg of total RNA were combined with 1 µL of 50 µM rRNA depletion oligos (about 380 nM each). Oligos were annealed to rRNA by incubating RNA at 98 °C for 2 minutes and then cooling to 22 °C (decreasing the temperature by 1 °C per second). After addition of 2 µL of 10× RNase H buffer at pH 8.3 [500 mM Tris-HCl, 750 mM KCl, 30 mM MgCl_2_, 100 mM DTT] and 1 µL (5 units) of thermostable RNase H (New England Biolabs), rRNA was digested at 50 °C for 30 minutes. Residual oligonucleotides were degraded by adding 24 µL of RNase-free water, 5 µL of 10× TURBO DNase Buffer, and 1 µL of TURBO DNase, and incubating at 37 °C for 30 minutes. rRNA-depleted RNA was purified by capture with 1.8× volumes of SPRI magnetic beads, three 70% ethanol washes, and 20 µL elution in RNase-free water.

### Ribosome-depleted RNA Sequencing Library Preparation

Reagents and primers used in RNA sequencing library preparation can be found in **Supplementary Table S10**. DNA sequencing libraries were generated from rRNA-depleted RNA using the NEBNext® Ultra™ II Directional RNA Library Prep Kit for Illumina largely according to manufacturer’s protocol. Broad steps include: RNA fragmentation, reverse transcription, dUTP-inclusive second strand synthesis, end prep and 3′ A-tailing, dUTP-including adaptor ligation, USER removal of dUTP nucleotides, and PCR amplification. We substituted our own dUTP-inclusive adapter mix that also included unique molecular identifiers and sequences amenable to our own multiplex barcoding strategy. We measured final library concentration by Qubit™ dsDNA HS Assay Kit and confirmed size distributions by D1000 HS Tape (TapeStation 4150 instrument, Agilent).

### Sequencing

Biological triplicate cDNA libraries were pooled and sequenced at low depth on an Illumina MiSeq for quality control, followed by high-depth sequencing on NextSeq and NovaSeq instruments in a 150 × 150 paired-end format. FASTQ files were demultiplexed on-instrument. Adapter/barcode and transposase-end sequences were removed in successive Cutadapt (**57**) passes. Fastp (**58**) performed quality trimming and extracted a 12-nt UMI from the 5′ end of each read. Paired reads were aligned without merging using HISAT2 (**59**) to the NCBI RefSeq GRCh38.p14 primary assembly (GCF_000001405.40), with RF strandedness. PCR duplicates were removed from paired reads using UMI-tools (**60**). FeatureCounts (**61**) assigned paired fragments to exons using the RefSeq primary GTF, grouped by gene_id and counted with reverse-stranded mode. Gene-level raw counts were analyzed with DESeq2 (**62**) using metadata-defined pairwise contrasts. Genes were retained for each comparison if they had at least 10 counts in at least two samples (this was critical for observation of genes like NMRAL2P that are entirely undetectable in cells without active NRF2).

### Plasmidsaurus RNA-Seq of gapmer and siRNA samples

DNase-treated total RNA from A549 and SKMES1 gapmer and siRNA experiments, including corresponding controls, were shipped on dry ice to Plasmidsaurus for 3′ end counting RNA-seq. Libraries were prepared using oligo(dT)-primed reverse transcription with UMI incorporation, followed by second-strand synthesis, tagmentation, amplification, and Illumina sequencing. Reads were quality filtered with fastp (**58**), aligned to the human reference genome using STAR (**63**), deduplicated using UMICollapse (**64**), and quantified using featureCounts (**61**) with strand-specific assignment to exons and 3′ UTRs, grouped by gene_id. Differential expression was analyzed by Plasmidsaurus using edgeR v4.0.16 (**65**) for A549 samples and the edgePython quasi-likelihood workflow for SKMES1 samples. Both analyses reported log_2_ fold changes, nominal P values, and FDR-adjusted P values with equivalent interpretations. Full differential-expression tables were downloaded as CSV files for each treatment-versus-control comparison and standardized to a common format. Contrast labels were oriented so that positive log_2_ fold changes indicated higher expression in the first named condition (gapmer or siRNA treatment), and results from both analyses were processed identically for downstream RRHO analysis using fold-change direction and nominal P values.

### RRHO2 analysis

DESeq2 contrast tables were oriented so that positive log_2_ fold change indicated higher expression in the first named condition (Cancer, Treatment, Mutation); contrasts with the opposite orientation were sign-reversed before analysis. For each pair of contrasts, genes with finite log_2_ fold-change and nominal P values in both tables were retained. Genes were ranked by signed significance, calculated as sign(log_2_ fold change) × −log_10_(P). RRHO2 (**66**) compared the ranked lists across rank cutoffs using its default step size and hypergeometric test, generating four-quadrant heatmaps for concordant and discordant changes. Heatmaps display −log_10_(P) without correction across the rank-cutoff grid. Plasmidsaurus edgeR contrast results were oriented similarly and ranked as sign(log_2_ fold change) × −log_10_(*P*), using the nominal *P* values from edgeR quasi-likelihood tests. For each analysis group, RRHO2 peak-overlap genes were collected separately by quadrant across all pairwise combinations of contrasts. Genes present in every pairwise peak list formed the group’s consensus signature, and overlaps among these signatures were summarized using Venn diagrams.

### TCGA survival analysis

TCGA-LUAD and TCGA-LUSC STAR-derived expression (star_tpm.tsv), somatic mutation (somaticmutation_wxs.tsv), and overall-survival (survival.tsv) datasets corresponding to GDC release 40.0, together with clinical (clinical.tsv) datasets corresponding to GDC release 41.0, were downloaded from the UCSC Xena GDC hub (**100**). Expression values were analyzed on the supplied log_2_(TPM + 1) scale. Analyses included primary tumors, retaining one sample per patient; patients lacking valid expression or survival data were excluded from survival analyses. Mutation-defined cohorts comprised patients with protein-altering *KEAP1* variants or *NFE2L2* variants within DLGex (aa 17-51) or the ETGE motif and its five-amino-acid flanks (aa 74-87). Patients carrying nonsynonymous *NFE2L2* variants of unknown significance outside these regions were excluded. Remaining patients were classified as having no reported qualifying mutation.

Separate expression-defined cohorts were established using the 10th percentile of NQO1 expression among mutation-positive patients within each cancer type: 7.669 for LUAD and 7.916 for LUSC. Patients were classified above or below these thresholds. Within each mutation- or NQO1-defined cohort, patients were ranked by NMRAL2P RNA expression and divided into upper and lower halves for LUAD or the bottom 25% and remaining 75% for the LUSC survival analyses. Expression ties at group boundaries were resolved by patient barcode. Overall survival was estimated using Kaplan-Meier curves (**67**) with 95% confidence intervals calculated from Greenwood’s variance using a log-log transformation. Survival distributions were compared using log-rank tests (**68**). Cross-cancer NMRAL2P expression plots included all primary-tumor patients with valid expression values, independently of mutation or survival data availability.

Cox proportional hazards models (**69**) evaluated NMRAL2P expression as a continuous predictor of overall survival, with hazard ratios expressed per one-standard-deviation increase in expression. Models were fitted separately for LUAD and LUSC and included NMRAL2P alone or with mutation category or NQO1 category, with additional adjustment for age, sex, and pathological stage. Clinical comparisons used complete cases without imputation; stage substages were combined into stages I-IV. Expression scaling was fixed using the original analysis population for each input dataset. Models were fitted using lifelines (**70**), and proportional hazards assumptions were assessed using tests based on Schoenfeld residuals (**71**). Reported P values were nominal and were not adjusted for multiple analyses.

### Gene set enrichment analysis

Preranked GSEA was performed using GSEApy (**72**) with differential expression results from DESeq2 or Plasmidsaurus edgeR/edgePython analyses. Genes were ranked by log_2_ fold change, oriented so that positive values indicated higher expression in the altered condition relative to its control. Genes lacking valid identifiers, finite fold changes, or nominal P values were excluded. Duplicate gene identifiers were resolved by retaining the entry with the largest absolute fold change, with nominal P value used to resolve ties. Exact fold-change ties were ordered using nominal P values, prioritizing stronger evidence toward the corresponding end of the ranked list. No differential expression significance threshold was applied before enrichment analysis.

Enrichment was evaluated against a combined 42-gene NRF2 core gene set (**73**) and selected KEGG signaling gene sets supplied in GMT format. Normalized enrichment scores, nominal P values, and FDR q values were obtained from GSEApy. Gene sets with FDR q ≤ 0.05 were considered significantly enriched. Leading-edge fractions were calculated as the number of leading-edge genes divided by the number of gene set members represented in the ranked list.

### Comparative sequence analysis

Genomic sequences were aligned using MAFFT (**74**) within Jalview (**75**) and manually curated to align homologous splice junctions and exon-intron boundaries. Exon and coding-region annotations were projected onto the resulting alignments.

Human *NMRAL1* and *NMRAL2P* genomic loci from GRCh38/hg38 were compared across their annotated gene regions. Gene structures and nucleotide conservation were visualized in shared alignment coordinates. Pairwise nucleotide identity was calculated across positions containing nongap nucleotides in both sequences.

Human-mouse comparisons were performed separately for *NMRAL1* and *NMRAL2P* using GRCh38/hg38 and GRCm39/mm39 genomic sequences. Conservation was evaluated across annotated exons, untranslated exonic regions, coding regions, introns, and flanking sequences. Identity excluding gaps was calculated as matches divided by matches plus mismatches; identity including indels was calculated as matches divided by all alignment positions containing a nucleotide in either species. Indel events were defined as contiguous runs of gaps in either sequence, and their frequencies and lengths were summarized by region.

NMRAL2P open reading frame conservation was additionally examined across human, chimpanzee, gorilla, rhesus macaque, and gray mouse lemur.

### Isoform analysis by full-length cDNA amplification

NMRAL2P isoforms were identified using the Lexogen TeloPrime Full-Length cDNA Amplification Kit V2. Total RNA was isolated, DNase-treated, and confirmed to have an RNA integrity number greater than 8. Full-length cDNA was generated according to the manufacturer’s instructions and then amplified using Q5 Hot Start High-Fidelity 2× Master Mix with the TeloPrime forward primer and an NMRAL2P-specific reverse primer (**Supplementary Tables S11 and S12**).

PCR conditions consisted of initial denaturation at 98 °C for 1 min; 20-30 cycles (depending on cell line) of denaturation at 98 °C for 10 s, annealing for 30 s at 64 °C, and extension at 72 °C for 90 s; followed by a final extension at 72 °C for 2 min. Amplified cDNA was purified using a 1× bead-to-sample volume ratio of Mag-Bind TotalPure NGS SPRI beads (Omega Biotek), with two washes in 80% ethanol and elution in 20 µL of RNase-free water. Amplicon sizes were assessed using an Agilent D5000 ScreenTape assay before submission to Plasmidsaurus for Oxford Nanopore long read sequencing.

Oxford Nanopore reads were processed using the Galaxy platform (**76**). Adapter sequences were removed using Porechop, and trimmed reads were aligned to the human reference genome assembly using Minimap2 (**77**) in splice-aware mode. Transcript structures were reconstructed and their abundances estimated using StringTie (**78**) with long-read settings and reference annotation. Transcripts corresponding to the NMRAL2P locus were identified from their genomic alignments and exon structures (**Supplemental Table S13**). GffRead (**79**) was used to extract transcript sequences from the resulting annotation and reference genome. Relative isoform abundance was calculated as the abundance assigned to each isoform divided by the total abundance of NMRAL2P isoforms in each sample.

### Cell Fractionation

To fractionate and isolate RNA, ∼85% confluent cells on a 10 cm dish were washed once with 1× PBS and then scraped into 350 µL of cold cell lysis buffer [40 mM Tris-HCl pH 8.0, 25 mM NaCl, 6 mM MgCl_2_, 1 mM CaCl_2_, 250 mM sucrose, 0.15% IGEPAL CA-630] supplemented with 800 units/mL of recombinant RNasin (Promega). 100 µL of the initial lysate was diluted with 150 µL of water and combined with 750 µL of TRIzol-LS: this is the whole cell fraction, stored at 4 °C until further processing. The remaining lysate was incubated on ice for 20 minutes and nuclei were pelleted for 5 minutes at 2, 300 × g at 4 °C. 250 µL of supernatant was carefully removed into a new tube and combined with 750 µL of TRIzol-LS: this is the cytoplasmic fraction, stored at 4 °C until further processing. The nuclear pellet was washed once with 300 µL of cold cell lysis buffer and pelleted again for 5 minutes at 2, 300 × g at 4 °C. The supernatant was discarded, and nuclei were lysed in 1 mL of TRIzol: this is the nuclear cell fraction, stored at 4 °C until processing. All fractions were processed simultaneously according to TRIzol manufacturer’s protocol, treated with DNase, and purified with SPRI beads as with total RNA preparations (see “RNA extraction and DNase treatment”). Fractionation reagents can be found in **Supplemental Table S14**.

### Reverse transcription and quantitative PCR (RT-qPCR)

cDNA was synthesized from 0.5 µg of RNA using the iScript gDNA Clear cDNA Synthesis Kit (Bio-Rad) according to the manufacturer’s instructions. Each sample included a reverse transcription reaction and a matched no reverse transcriptase (no RT) control. cDNA was diluted fivefold in molecular-grade water. Half-volume qPCR reactions contained diluted cDNA, 1 µM each forward and reverse primer (**Supplementary Table S12**), and 1× PowerUp SYBR Green Master Mix (Thermo Fisher Scientific). Reverse-transcribed samples were analyzed in technical triplicate, and no RT controls were analyzed in a single well per primer pair. Reactions were run on a Bio-Rad CFX Opus instrument with an initial incubation at 95 °C for 3 min, followed by 40 cycles of denaturation at 95 °C for 15 s, annealing at 65 °C for 30 s, and extension at 72 °C for 40 s. No RT controls were required to amplify at least eight quantification cycles (C_q_) later than the corresponding reverse transcribed samples. RT-qPCR reagents are listed in **Supplemental Table S15**.

Target RNA measurements were normalized to 18S rRNA using ΔC_q_ = C_q_(target) − C_q_(18S), calculated from the mean technical replicate C_q_ values. Fold changes following gapmer or siRNA treatment were calculated relative to the corresponding control using the 2^⁻ΔΔCq^ method. For nuclear-cytoplasmic localization analyses, normalized relative quantities were calculated separately for each fraction as 2^⁻ΔCq^. Each fraction’s relative quantity was divided by the sum of the nuclear and cytoplasmic relative quantities, yielding proportions that summed to 1. These values represent 18S-normalized relative proportions rather than absolute fractions of cellular RNA; localization was interpreted by comparison with reference RNAs of established nuclear or cytoplasmic localization.

### NMRAL2P ORFp structure prediction and comparison

The structure of the major NMRAL2P open reading frame product (ORFp) was predicted using AlphaFold3. Predicted models were structurally aligned to the NADPH-bound and unbound NMRAL1 subunits in the crystallographic asymmetric dimer (PDB: 2EXX) to compare their folds and spatial relationships to the NADPH-binding pocket and dimerization interface. AlphaFold-Multimer was used to predict NMRAL1-NMRAL2P ORFp, NMRAL1-ASS1, and NMRAL2P ORFp-ASS1 complexes. Predicted structures were colored by model confidence, and surface representations of structurally aligned NMRAL1 or NMRAL2P ORFp were overlaid for comparison.

### In-cell chemical probing with 5NIA

A549, SKMES1, and H1299 (H1299 cells were treated with 100 nM CDDO-Me 24 hr prior to in-cell chemical probing) cells were grown to 70–80% confluence in 10-cm dishes and washed twice with 1× PBS. Each dish received 4.5 mL of 1× PBS followed by 500 µL of freshly prepared 250 mM 5NIA (Sigma) in DMSO, yielding a final concentration of 25 mM 5NIA and 10% DMSO. Vehicle-control dishes received 500 µL of DMSO. Cells were incubated for 10 min at 37 °C, after which the solution was aspirated and cells were washed once with 1× PBS. RNA was isolated as described in “RNA extraction and DNase treatment”. Chemical probing reagents are listed in **Supplementary Table S16**.

### Selective 2′-hydroxyl acylation analyzed by primer extension and mutational profiling (SHAPE-MaP)

DNase-treated RNA from 5NIA-treated and DMSO-control samples were reverse transcribed under MaP conditions using SuperScript II (SSII). RNA (5 µg in 7 µL RNase-free water) was combined with 2 µL of 10 mM dNTPs and 1 µL of either random DNA nonamers (200 ng/µL) or 2 µM gene-specific RT primer in a total volume of 10 µL. The RNA mixture was denatured at 70 °C for 5 min and then cooled at 4 °C for 2 min. Each sample received 9 µL of freshly prepared MaP buffer containing 2 µL of 10× buffer [500 mM Tris-HCl (pH 8.0), 750 mM KCl, 100 mM DTT], 4 µL of 5 M betaine, and 3 µL of 40 mM MnCl₂, with MnCl₂ added immediately before use. Samples were incubated at 25 °C for 2 min before addition of 1 µL (200 units) SSII. Reverse transcription proceeded at 25 °C for 10 min and 42 °C for 90 min, followed by 10 cycles of 50 °C for 2 min and 42 °C for 2 min, and enzyme inactivation at 70 °C for 10 min. cDNA was buffer exchanged using MicroSpin G-50 columns (Cytiva).

Sequencing libraries were generated by two successive PCR amplifications using Q5 Hot Start High-Fidelity 2× Master Mix. Target-specific amplification (NMRAL2P was split into multiple overlapping amplicons ∼400 nt each depending on isoform) was performed in 25 µL reactions containing 1-3 µL of buffer-exchanged cDNA, 1.25 µL each of 10 µM forward and reverse primers (**Supplementary Table S12**), and 12.5 µL of 2× master mix. Reactions were incubated at 98 °C for 1 min, followed by 15 cycles of 98 °C for 10 s, primer-specific annealing for 30 s, and 72 °C for 30 s, with a final extension at 72 °C for 2 min. Products were purified using a 1× SPRI bead cleanup, washed twice with 80% ethanol, and eluted in 15 µL of molecular-grade water. DNA concentrations and fragment sizes were assessed using the Qubit dsDNA HS and Agilent D1000 HS ScreenTape, respectively.

Indexing PCR was performed in 50 µL reactions containing 2 ng of purified target-specific PCR product, 5 µL each of 5 µM forward and reverse indexing primers (**Supplementary Table S10**), and 25 µL of Q5 master mix. Cycling conditions were 98 °C for 1 min; 10 cycles of 98 °C for 10 s, 67 °C for 30 s, and 72 °C for 30 s; and a final extension at 72 °C for 2 min. Indexed libraries were purified using a 0.8× SPRI bead cleanup, washed twice with 80% ethanol, and eluted in 15 µL of molecular-grade water. Library concentrations and sizes were assessed using Qubit dsDNA HS and Agilent D1000 HS ScreenTape. Pooled libraries were sequenced on an Illumina MiSeq instrument to generate 2 × 250-bp paired-end reads.

### SHAPE-MaP data processing and RNA secondary structure analysis

Sequencing reads from 5NIA-treated and matched DMSO-control samples were processed using Shapemapper2 (**80**) against the corresponding NMRAL2P reference sequences. Per-nucleotide mutation rates were calculated, background-corrected using the DMSO controls, and normalized to generate SHAPE reactivity profiles. Isoform-specific amplicons were analyzed separately to generate reactivity profiles for the major NMRAL2P isoforms.

Sequence-only folding analyses were performed for each isoform using the “partition” function within RNAstructure (**81, 82**). Base-pairing probabilities were extracted using ProbabilityPlot and visualized using ArcPlot (**83**). These analyses used nucleotide sequence alone, without SHAPE-derived restraints, and produced arc diagrams representing pairing probabilities across the predicted secondary structure ensemble.

### Treatment/transfection of siRNA and gapmers

Lyophilized gapmers were reconstituted in a 1× IDTE buffer at pH 7.5 (10 mM Tris-HCl pH 7.5 and 0.1 mM EDTA), aliquoted, and stored at −20 °C. NRF2 siRNA (Thermo Fisher Scientific) were diluted to 20 mM working stocks and stored at −70 °C (**Supplementary Table S17**).

Before transfection, A549 and SKMES1 cells were seeded into 6-well plates at 5×10^5^ and 7×10^5^ cells per well, respectively. 24 hr after seeding, cells were transfected with either 25 nM gapmer or 20 nM siRNA final concentration using Lipofectamine 2000 per the manufacturer protocol. For each well, 7.5 µL of Lipofectamine 2000 and the appropriate amount of gapmer or siRNA were diluted separately in RPMI medium to a final volume of 150 µL each. After incubation for 5 min at room temperature, the diluted Lipofectamine 2000 and oligonucleotide solutions were combined (300 µL total) and incubated for an additional 10-15 min at room temperature. Mixtures were added dropwise to appropriate wells. Cells were processed 24 hr post transfection for RNA harvest, proliferation assays, and/or RT-qPCR.

### Proliferation Assays

24 hr after gapmer or siRNA transfection, cells were washed once with 1× PBS and split with 0.25% trypsin. Cells were counted and A549 and SKMES1 cells were seeded 2, 500 and 7, 000 cells in 100 μL volume, respectively per well of a 96-well plate. Plates were incubated and imaged every 2 hours for 4 days in an IncuCyte S3 (Sartorius). Analysis was performed using the IncuCyte adherent cell-by-cell analyses (individual cell line specific trained analyses were used for each respective cell line) and proliferation rates were calculated for each cell line by normalizing cell count to time zero. For proliferation assays, 4 biological replicates were performed for A549, and 3 biological replicates were performed for SKMES1 cells.

Cells not used for plating were washed twice with 1× PBS, pelleted at 300 x g for 5 minutes, and harvested for RNA with TRIzol ( see RNA extraction and DNase treatment) and used to validate knockdown efficiency using RT-qPCR.

### Cisplatin Sensitivity Assays

Cisplatin (Thermo Fisher Scientific) was resuspended in 150 mM NaCl_2_ with gentle heating to 37 °C and occasional vortexing, and stored at −20 °C until use within 1 month of being resuspended. Cisplatin treatment followed proliferation assay workflow: post-gapmer or -siRNA treatment, A549 and SKMES1 cells were seeded at 2, 500 and 7, 000 cells in 90 μL volume, respectively per well in 96-well plate. Cells were left to adhere to the plate for three to four hours before adding 10 μL of cisplatin or saline diluted in respective cell line media to achieve the desired final cisplatin dose. 96 hr post-treatment, cell number relative to time 0 was measured by IncuCyte adherent cell-by-cell analyses (as in our proliferation assay). Endpoint counts were plotted against cisplatin concentration to derive IC_50_ (half-maximal inhibitory concentrations). For cisplatin sensitivity assays, 4 biological replicates were performed for A549 cells, and 3 biological replicates were performed for SKMES1 cells.

### Functional over-representation analysis

The RRHO2-derived gene signature concordantly downregulated by NMRAL2P gapmers in A549 and SKMES1 cells, excluding genes belonging to the shared NRF2-siRNA signature, was analyzed using the g:GOSt web interface from g:Profiler (**84**). Human KEGG pathways were tested using an unordered query and a custom background comprising genes tested in all eight gapmer and siRNA comparisons. Following identifier mapping, the analysis included 824 query genes and an effective background of 13, 631 genes. Enrichment was assessed using cumulative hypergeometric tests with g:SCS multiple-testing correction and an adjusted P-value threshold of 0.05. Results were downloaded with gene intersections, and fold enrichment was calculated as observed overlap divided by expected overlap using the exported query, pathway, and background sizes.

### Pathway activity scoring

Gene-count matrices were processed separately for A549 and SKMES1. Counts sharing a gene symbol were summed, low-expression genes were removed using edgeR filterByExpr with treatment group specified, and libraries were normalized using the trimmed mean of M values (TMM). Expression was converted to log_2_ counts per million using edgeR with a prior count of 2. All 24 samples within each cell line, including untreated and transfection-reagent controls, contributed to normalization and score reference distributions.

The lung MEK score was calculated as the mean normalized log_2_ expression of *DUSP4*, *DUSP6*, *ETV4*, *ETV5*, *PHLDA1*, and *SPRY2*, adapting this six-gene signal (**85**) to RNA-seq data. The MAPK pathway activity score (MPAS (**86**)) included *PHLDA1*, *SPRY2*, *SPRY4*, *DUSP4*, *DUSP6*, *CCND1*, *EPHA2*, *EPHA4*, *ETV4*, and *ETV5*. Expression of each gene was standardized using its mean and sample standard deviation across the 24 samples within its cell line; MPAS was calculated as the sum of these gene-level z scores divided by √10. All signature genes were required to pass expression filtering.

PROGENy pathway scores were calculated from TMM-normalized log_2_ counts per million using the human model with the top 100 response genes per pathway. Each score was calculated as the sum of available genes’ expression values multiplied by their signed model weights. All 14 pathways were scored without standardizing the resulting scores or performing permutation-based significance testing.

### Normalization of pathway score shifts

For lung MEK, MPAS, and PROGENy MAPK scores, shifts were calculated as the mean treatment score minus the mean corresponding control score, divided by the sample standard deviation of that score across all 24 samples within the same cell line, including untreated and transfection-reagent controls. Gapmers were compared with scramble controls and NFE2L2 siRNA with control siRNA. The same reference standard deviation was used for all treatment comparisons within each score and cell line. Multi-pathway PROGENy shifts were calculated identically for each pathway. Positive and negative shifts indicate increased and decreased scores relative to the corresponding control, respectively.

## RESULTS

### Disrupting KEAP1-NRF2 interaction activates NRF2 antioxidant signaling in lung and lung cancer cells

Under basal conditions, KEAP1 promotes NRF2 ubiquitination and proteasomal degradation, whereas oxidative stress inhibits this process, allowing NRF2 to accumulate and activate an antioxidant gene expression response (**Fig. 1A**). Analysis of Cancer Dependency Map data (**54, 87, 88**) identified *NFE2L2*, which encodes NRF2 protein, as the most significantly enriched dependency in NSCLC relative to other cancer cell lines (**Fig. 1B**): specifically, 38.8% of NSCLC lines were classified as *NFE2L2*-dependent (**Fig. 1C**). This dependency extended beyond NSCLC: 18.1% of head and neck squamous cell carcinoma lines and 13.7% of all cancer cell lines analyzed were also classified as *NFE2L2*-dependent (**Fig. 1C** and **Supplementary Fig. S1A**). These findings motivated our investigation of NRF2-dependent transcriptional programs as potential therapeutic vulnerabilities in NSCLC.

First, we confirmed the activation status of the NRF2 pathway across normal lung and NSCLC cell lines. We found that cells without mutations in the NRF2 pathway – SV40-immortalized but otherwise normal human bronchial epithelial cells 16HBE14o-(HBE), adenocarcinoma H1299, and squamous cell carcinoma CALU1 – expressed low levels of NRF2 protein and NRF2-activated target proteins including NQO1, SLC7A11, SQSTM1, and TXNRD1 (**Fig. 1D**). We confirmed that NRF2 pathway activity in these NRF2-inactive cells could be dramatically stimulated by treatment with the small molecule bardoxolone methyl (methyl 2-cyano-3, 12-dioxooleana-1, 9(11)-dien-28-oate or CDDO-Me) (**Fig. 1D**). These CDDO-Me-activated cells represented cells with an acute short-term NRF2 response: even within 24 hours, we observed strong activation of protein expression from NRF2 target genes. To observe any differences in gene expression after chronic long-term NRF2 activation, we generated H1299 cell lines stably expressing NRF2 with a targeted point mutation (NRF2^E79Q^) or including Cas9-sgRNA-driven frameshift loss-of-function indels in the *KEAP1* gene (sgKEAP1): in both cases KEAP1-dependent NRF2 turnover is inhibited and NRF2 is constitutively active (**Fig. 1E**). We also confirmed basal NRF2 protein abundance and expression of NRF2 target proteins in the lung adenocarcinoma cell lines A549 and H2228 and the lung squamous cell carcinoma lines H520 and SKMES1 (**Fig. 1F**). A549 harbors a loss-of-function *KEAP1* p.G333C mutation (**34, 89**) and *KEAP1* promoter hypermethylation (**90**), while H2228 harbors the activating *NFE2L2* p.G31A mutation (**91**). Elevated basal NRF2 protein abundance has been reported in both H520 and SKMES1 cells, together with an increased *NFE2L2/KEAP1* copy-number ratio (**92, 93**). These cell lines (A549, H2228, H520, SKMES1), having higher basal levels of NRF2, are consequently less sensitive to further activation by CDDO-Me (**Supplementary Fig. S1B**). While there is cell-type specific variation among NRF2 target gene expression, we classified A549, H2228, H520, and SKMES1 cells as NRF2-active cancer cell lines for subsequent analyses.

### Long noncoding RNA NMRAL2P is among a small group of transcripts significantly activated upon all methods of NRF2 activation in lung and lung cancer cells

After confirmation of NRF2 activity, we set out to determine NRF2-dependent genes, measuring activated transcripts by a standard strand-specific RNA sequencing approach (**Fig. 2A**). To enable recovery of non-adenylated transcripts in our data, we performed RNase H-based rRNA depletion (**94, 95**) instead of poly(A) selection. To identify a true core set of NRF2-dependent genes, we looked at the overlap of three distinct categories of NRF2 activation, focusing on genes concordantly upregulated in every category (**Fig. 2B**). 1) In our “Cancer” group, we included genes upregulated in NRF2-active NSCLC cells (A549, H2228, SKMES1, and H520) compared to normal lung HBE cells. 2) Our “Treatment” category included genes upregulated when each NRF2-inactive cell line (HBE, H1299, CALU1) was treated with CDDO-Me for 24 hours. 3) Lastly, our “Mutation” group included genes upregulated in an NRF2-inactive NSCLC line (H1299) when the NRF2 pathway was internationally altered with stable expression of hyper stable NRF2^E79Q^ or CRISPR-guided removal of KEAP1 protein expression.

**Figure 2:**
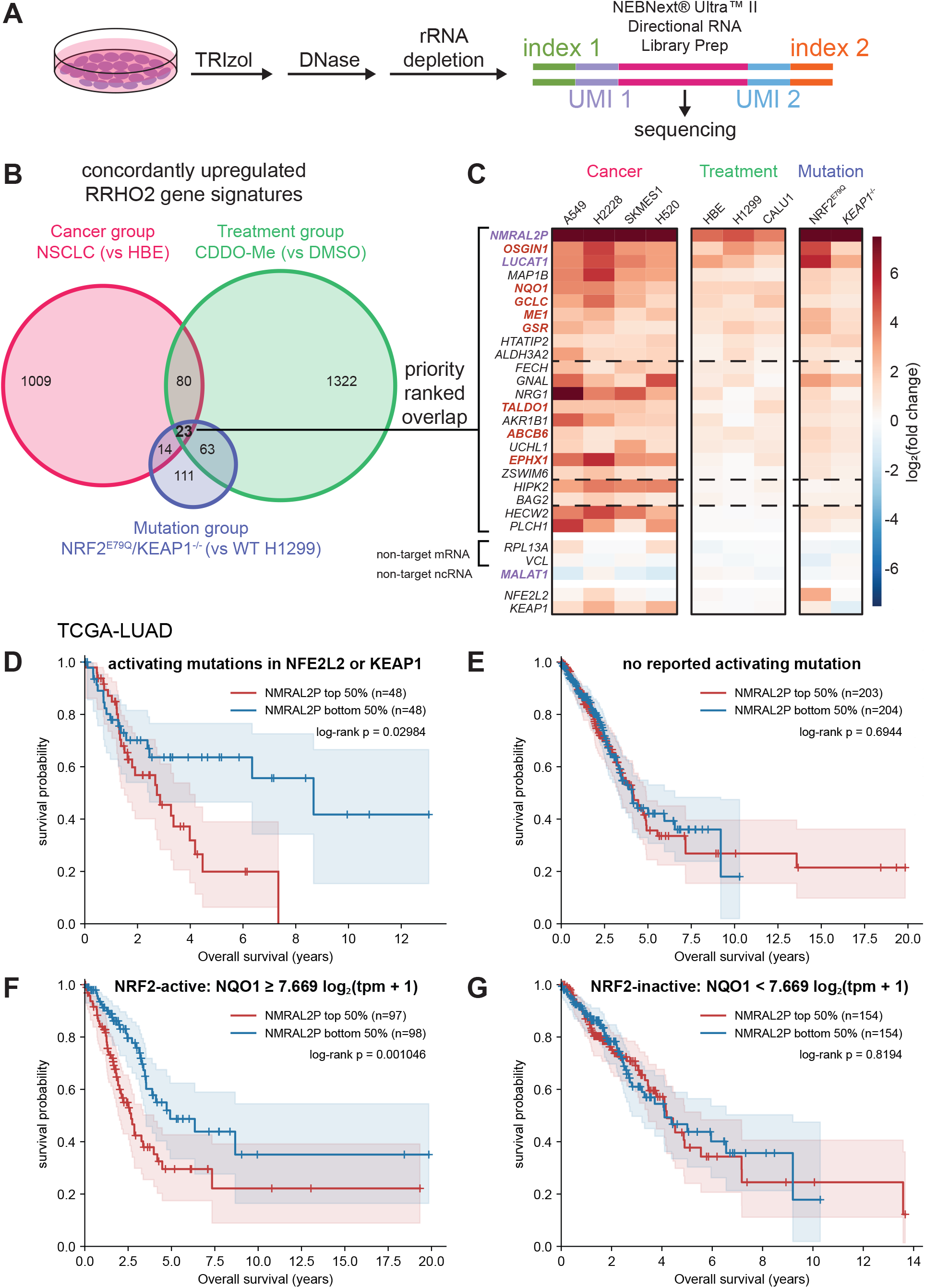
Long noncoding RNA NMRAL2P is expressed upon NRF2 activation in lung and lung cancer cells and is a potential liability for survival in lung adenocarcinoma patients. (**A**) Library preparation schematic for RNA-seq of lung and lung cancer cell lines. (**B**) Venn diagram representing intersection of indicated groups’ rank-rank hypergeometric overlaps. (**C**) Heat map of log_2_(fold-change) for each indicated cell comparison. Overlapping NRF2 activated genes from A are ordered by priority ranking. Genes above dotted lines achieved BH-adjusted P < 0.05 in 9/9, 8/9, and 7/9 samples, respectively. Non-target control genes, *KEAP1*, and *NFE2L2* are also included. Core NRF2 target gene names are colored red while noncoding RNA gene names are colored purple. Kaplan-Meier survival plots for TCGA lung adenocarcinoma patients with (**D**) or without (**E**) NRF2-activating mutations and with (**F**) or without (**G**) NRF2 activity based on NQO1 RNA expression are shown. A “+” mark represents a censored observation, and shaded regions include 95% confidence intervals.

To identify genes consistently activated across these diverse models, we employed rank-rank hypergeometric overlap (RRHO) analysis within each group, which identifies concordant transcriptional changes between datasets without imposing arbitrary differential expression thresholds (**66, 96**). Pairwise comparisons within both the Cancer and Treatment groups revealed extensive overlap among concordantly upregulated and downregulated genes, whereas comparisons within the Mutation group identified a smaller set of concordantly upregulated genes, with little evidence of shared downregulation (**Supplementary Fig. S2**). These distinct patterns suggest that the Cancer and Treatment groups capture broad transcriptional changes associated with cancer cell states and pharmacological treatment, respectively, whereas genetic activation of NRF2 produces a more restricted, predominantly activating transcriptional response consistent with the targeted perturbation of a single transcriptional regulatory pathway. Still, every cell line and condition tested showed a significant enrichment of upregulated NRF2 target gene expression (**Supplementary Fig. S3**)

For each category, we defined a stringent set of concordantly upregulated genes by retaining only genes present in the enrichment peaks corresponding to upregulation in both datasets in every pairwise RRHO comparison. This yielded three category-specific sets of consistently upregulated genes, which were subsequently intersected to identify a core transcriptional signature shared across Cancer, Treatment, and Mutation groups (**Fig. 2B**). We ranked the 23 genes at the intersection, prioritizing genes with the greatest fraction of comparisons exhibiting positive log_2_(fold changes) and statistically significant differential expression, followed by the median magnitude of upregulation and adjusted p-value (**Fig. 2C**). Among the list, 8 genes part of a core NRF2 activation program (**73**) clustered at the top of our ranking. Interestingly, at the top of the list (position 1 and 3) were two long noncoding RNAs (lncRNAs), NMRAL2P and LUCAT1. Based on its unique enrichment across our datasets, we decided to investigate the *NMRAL2P* gene further.

NMRAL2P (also known as LOC344887) is an NRF2-responsive lncRNA whose direct transcriptional activation by NRF2 has been demonstrated in colon cancer cells and lung cell models of pulmonary fibrosis (**50, 51**). Reported downstream functions include promoting NQO1 induction in sulforaphane-treated colon cancer cells and suppressing profibrotic gene expression in lung cells. In head and neck cancer cells, NMRAL2P has also been reported to associate with NRF2 protein and enhance NRF2-mediated *GPX2* transcription (**52**). NMRAL2P expression is elevated in NSCLC tumors relative to nonmalignant lung tissue (**97, 98**), and higher tumor expression of NMRAL2P was reported to predict shorter overall survival in a cohort of 173 patients who underwent NSCLC resection at two hospitals in Shandong, China (**53**).

To assess the clinical relevance of the NRF2-NMRAL2P relationship, we analyzed patient data from the The Cancer Genome Atlas (TCGA) lung adenocarcinoma (LUAD) and lung squamous cell carcinoma (LUSC) cohorts (**16, 99**), accessed through UCSC Xena (**100**). First, we separated out patient samples with potentially NRF2 activating mutations from the LUAD cohort: either nonsynonymous alterations to *KEAP1* or nonsynonymous alterations corresponding to the KEAP1-interacting regions of NRF2 protein (DLGex residues 17-51 and ETGE-adjacent residues 74-87). Within both mutant and non-mutant cohorts, we split samples into top and bottom 50th percentiles by *NMRAL2P* expression. In the mutant cohort, higher levels of NMRAL2P were associated with significantly decreased survival (**Fig. 2D**), while there was no significant difference in survival correlated with NMRAL2P in the non-mutant cohort (**Fig. 2E**). Because missense mutations in *NFE2L2* and *KEAP1* do not represent the only mechanism of NRF2 activation, we performed the same analysis after creating “NRF2-active” and “NRF2-inactive” cohorts based on the expression of NRF2 target gene NQO1, under the assumption that cells achieving NQO1 expression equivalent to that of NRF2 mutant cells would result from NRF2 activity. We chose the threshold for NRF2-active NQO1 expression based on the 10th percentile of NQO1 expression in our mutant cohort (**Supplementary Fig. S4A** and **S4B**). In this more comprehensive “NRF2-active” cohort, patients with higher NMRAL2P expression had significantly worse overall survival rates compared to patients with lower NMRAL2P expression (**Fig. 2F**), while no difference was observed in the NRF2-inactive cohort (**Fig. 2G**). These observations suggest that NMRAL2P may be a therapeutic vulnerability in lung adenocarcinoma patients.

We performed a similar analysis for lung squamous cell carcinoma samples in TCGA (LUSC) but found no significant difference in survival in any cohort (**Supplementary Fig. S5A-F**), even when adjusting the cohort split between bottom 25th percentile and top 75th percentile to accommodate the order of magnitude increase in NMRAL2P expression in LUSC patient tumors (**Supplementary Fig. S5G**). Lung squamous carcinoma insensitivity to differences in NMRAL2P expression may be a consequence of a common LUSC chromosomal amplification of 3q26-3q28 (**101**), which includes the *NMRAL2P* gene. Cox regression (**69, 70**) treating NMRAL2P expression as a continuous variable likewise revealed no significant association with overall survival in LUSC, whereas the association in LUAD persisted after adjustment for age, sex, stage, and NQO1 expression category (**Supplementary Fig. S5H**). Whether NMRAL2P is of clinical relevance in LUSC requires testing by other means.

### NMRAL2P is a NRF2-dependent pseudogene-derived transcript retained in the nucleus of NSCLC cells

Very little is known about the *NMRAL2P* locus beyond few reports linking it to NRF2 transcriptional activity (**50–52**), so we decided to re-evaluate its origin and annotation through a series of genome alignments. *NMRAL2P* is annotated as a transcribed unprocessed pseudogene, meaning that RNA expression from a duplicated genetic locus is detectable but has lost coding potential for the original functional protein. *NMRAL2P* is syntenic throughout mammals, and in humans the *NMRAL2P* locus maintains 60.1% nucleotide identity with its duplication origin *NMRAL1* (protein product NMRAL1/HSCARG), including shared intron-exon distribution and 62.4% nucleotide identity to the original open reading frame (**Fig. 3A**). While the original ORF retained in *NMRAL2P* is generally conserved across primates (**Supplementary Fig. S6**), a human-specific mutation leaves a stop codon after 39 amino acids (**Fig. 3B**). A downstream, in-frame partial ORF is retained after the stop codon, but structural modeling predicts that its putative translation product would lack the native architecture required for NADPH binding and argininosuccinate synthase 1 inhibition (**Supplementary Fig. S7**), two linked functions of NMRAL1 (**102, 103**). The stop codon lies 88 nt upstream of the nearest downstream exon-exon junction, exceeding the canonical 50-55 nt boundary associated with exon-junction-complex-dependent nonsense-mediated decay (NMD) and predicting NMD susceptibility if this ORF is translated (**104, 105**). Interestingly, the level of conservation between mouse *Nmral2p* and human *NMRAL2P* genomic sequence is similar to that of mouse *Nmral1* and human *NMRAL1*, meaning that both elements are under equivalent selective pressure (**Supplementary Fig. S8**). While the coding sequence of *NMRAL1* is nearly completely conserved, the original ORF of mouse *Nmral2p* has accrued many indels that eliminate its coding potential. We did not find evidence of *NMRAL2P* transcription in other primates from existing databases or in tissues of mice with active NRF2 (**106**), suggesting that NRF2-dependent transcription of *NMRAL2P* could be a human-specific phenomenon.

**Figure 3:**
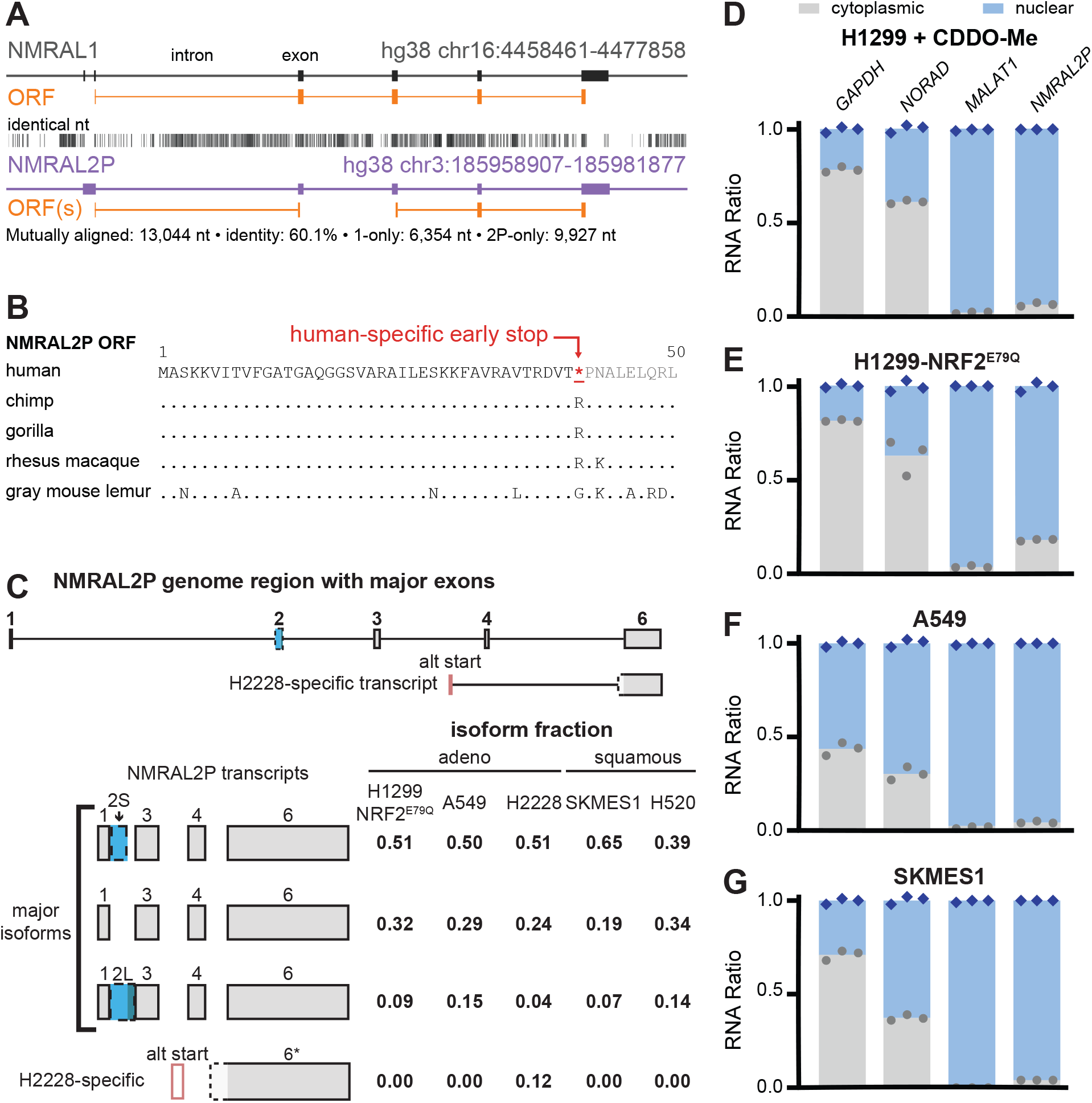
NMRAL2P is a NRF2-dependent pseudogene-derived transcript retained in the nucleus of NSCLC cells. (**A**) Alignment of human genes *NMRAL1* and *NMRAL2P*, with introns, exons, and open reading frames (ORF) labeled. Genome coordinates are displayed along with alignment information. (**B**) Conservation of the first 50 amino acids of the first NMRAL2P ORF across indicated primates. A human-specific early termination codon is highlighted in red. (**C**) Relative positions of major identified exons across the NMRAL2P genome region are indicated, along with positions for H2228-specific transcript (top). Exon schematics for each isoform along with their relative fraction of total NMRAL2P RNA from each indicated cell line (bottom). The RNA ratio of cytoplasmic to nuclear localization of NMRAL2P is compared against mRNA and lncRNA controls in (**D**) CDDO-treated H1299 cells, (**E**) H1299-NRF2^E79Q^ cells, (**F**) A549 cells, and (**G**) SKMES1 cells.

We investigated what, if any, isoforms of NMRAL2P transcripts are routinely expressed across NSCLC (**Fig. 3C**). Based on our RNA sequencing, we observed that the conserved exons of NMRAL2P were generally evenly expressed, but we were interested in their distribution across transcripts and whether the same transcripts were expressed across cell lines. To measure isoform distributions, we performed a cap-dependent ligation from NMRAL2P-specific RT-qPCR, adding sequencing adapters exclusively to full length cDNAs including from the 5′ cap to the end of the final exon (exon 6). All cDNAs were PCR amplified and subjected to long read sequencing, providing semi-quantitative isoform distributions of full length NMRAL2P transcripts. Among H1299-NRF2^E79Q^, A549, H2228, SKMES1, and H520 cells, a common pattern of 3 major isoforms emerged, varying only by whether exon 2 was included and which 5′ splice donor site was used (**Fig. 3C**). We also observed a variety of alternative splicing events that combined represent < 10% of observed transcripts, which we largely attributed to incomplete intermediates (**Supplementary Fig. S9**). H2228 cells also expressed an alternative isoform at appreciable levels (12% of transcripts) employing an alternative transcription start site and unique splice donor-acceptor pair (**Fig. 3C**). Interestingly, the three major isoforms retain one or both fragmented ORFs.

As the potential for generating translation products remained, we determined whether NMRAL2P transcripts persisted in the cytoplasm across NRF2-active NSCLC cell lines (**Fig. 3D-G**). We compared the amount of NMRAL2P RNA in the nucleus and cytoplasm by RT-qPCR after cellular fractionation, using levels of an mRNA (*GAPDH*), a cytoplasmic lncRNA (*NORAD*), and a nuclear lncRNA (*MALAT1*) as controls. In every NRF2-active condition – CDDO-Me-treated H1299 cells (**Fig. 3D**), H1299-NRF2^E79Q^ with activating NRF2 mutations (**Fig. 3E),** and in LUAD and LUSC cell lines (A549 and SKMES1) with intrinsically active NRF2 (**Fig. 3F-G**) – NMRAL2P transcripts were explicitly localized to the nucleus. While we did not determine the mechanism of nuclear localization (NMD or otherwise), the absence of a significant fraction of NMRAL2P transcripts in the cytoplasm supports the model of NMRAL2P remaining untranslated and with function restricted to the nucleus.

### In-cell chemical probing defines gapmer accessible sites in NMRAL2P

After defining the isoform distribution and localization of *NMRAL2P* transcripts, we sought to develop tools for modulating their expression. Given the predominantly nuclear localization of NMRAL2P, we used antisense gapmer oligonucleotides that recruit RNase H1 to cleave the target RNA (**107**), as this approach can achieve more efficient depletion of nuclear lncRNAs than RNA interference (**108**).

To identify flexible, protein-free regions of NMRAL2P accessible to gapmer binding, we chemically probed H1299-NRF2^E79Q^ and A549 cells by selective 2′-hydroxyl acylation analyzed by primer extension and mutational profiling (SHAPE-MaP) (**109**) using the more cell-amenable reagent 5-nitroisatoic anhydride (5NIA) (**110**). SHAPE-MaP identifies conformationally flexible nucleotides within RNA at single-nucleotide resolution: SHAPE adduct-induced “mutations” during specialized reverse transcription are measured by sequencing (**Fig. 4A**). For nucleotides to be reactive to 5NIA inside cells, they must reside within regions that are both unpaired and unbound, revealing regions available for base-pairing with gapmers. We mapped the accessibility of NMRAL2P major isoforms in both H1299-NRF2^E79Q^ and A549 cells (**Fig. 4B, 4C**, and **Supplementary Fig. S10A-D**). In general, average SHAPE reactivity of all isoforms between cell lines were very similar, suggesting that RNA structure and protein interaction networks are largely shared between the two NSCLC lines. Across all isoforms, SHAPE reactivity is generally lower across the 5′ segment of NMRAL2P including exons 1-4, while the shared terminal exon is more broadly SHAPE-reactive. The terminal exon is similarly reactive between LUAD and LUSC cell lines (**Supplementary Fig. S11A** and **S11B**).

**Figure 4:**
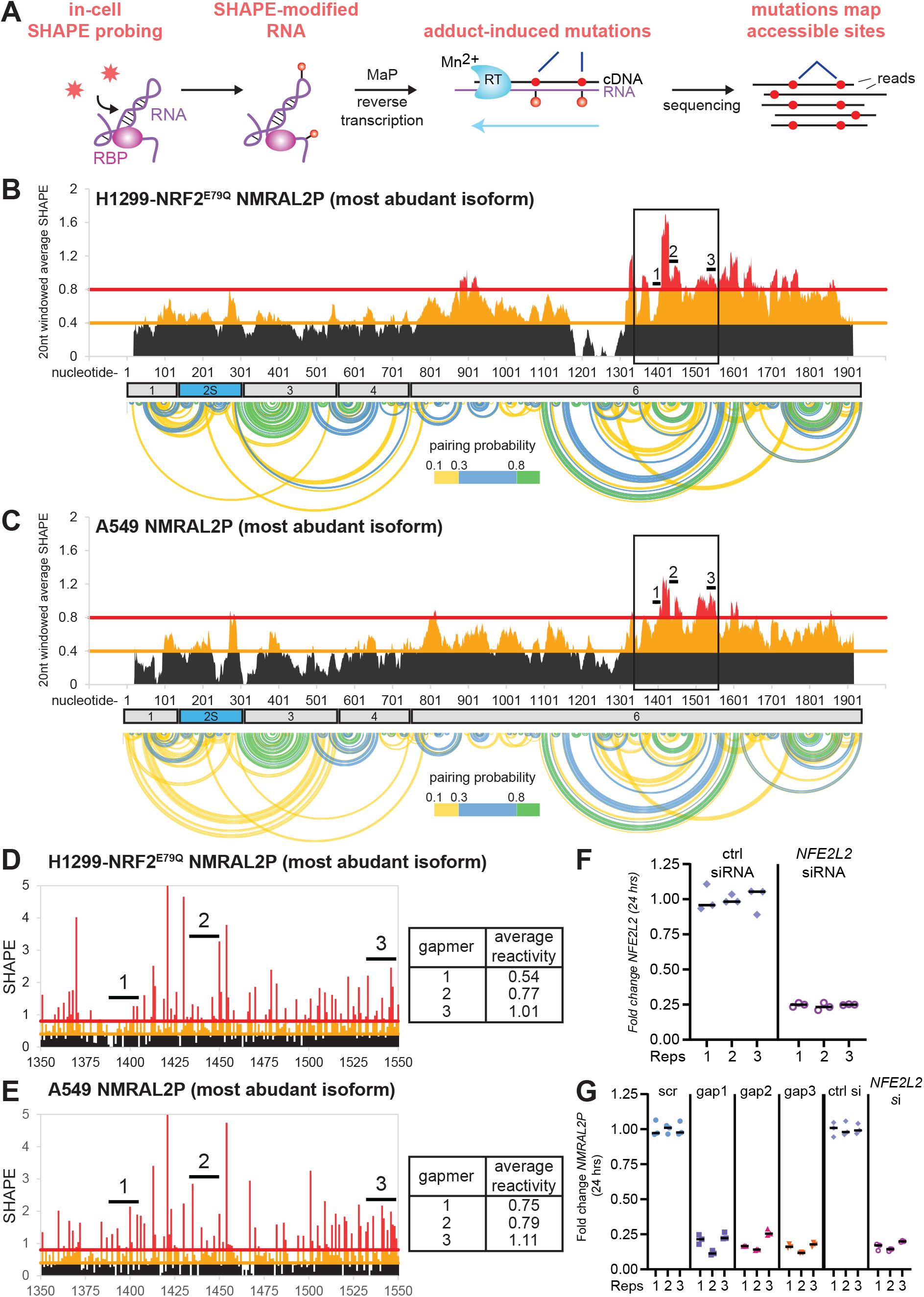
In-cell chemical probing defines gapmer accessible sites in NMRAL2P. (**A**) Schematic of in-cell SHAPE-MaP. Windowed average in-cell normalized SHAPE reactivity for the major NMRAL2P isoform from (**B**) H1299-NRF2^E79Q^ and (**C**) A549 cells. Exon locations and sequence-only informed pairing probability are aligned to the data. Black boxes highlight zoomed-in regions represented in (**D**) and (**E**), where nucleotide-resolution normalized SHAPE reactivity is plotted (nucleotide 1421 reactivity > 5). Average SHAPE reactivity in each gapmer target is displayed in tables. The fold change of (**F**) *NFE2L2* mRNA in A549 cells when exposed to indicated siRNAs and (**G**) NMRAL2P ncRNA in A549 cells when exposed to indicated gapmers and siRNAs. ctrl = control, si = short interfering RNAs, scr = scrambled gapmer (from gapmer 2 sequence), gap = gapmer ASO.

We decided to focus gapmer design against the SHAPE-reactive 3′ segment. We evaluated potential secondary structure through calculation of a nearest-neighbor informed partition function based on sequence alone (**Fig. 4B**, and **4C**). One 250 nt segment, retained in all major isoforms, maintained the highest average SHAPE reactivity and fewer well-determined potential secondary structures (fewer high probability pairings and significant alternative pairings). We designed three 16-nt antisense gapmers, each comprising a central 10-nt DNA gap flanked by three locked nucleic acid (LNA) residues at each end, with phosphorothioate linkages throughout. This design combines the enhanced RNA-binding affinity of LNA flanks and increased nuclease resistance of phosphorothioate linkages with a DNA gap that supports RNase H1-mediated target cleavage (**107, 111**). Gapmer target sequences lie outside of NMRAL1 conserved sequences. While all gapmers resided in high average SHAPE regions, the average SHAPE reactivity of each gapmer 16 nt target region varied between 0.5-1.0 (**Fig. 4D** and **4E**).

We then confirmed NMRAL2P knockdown from each gapmer. As a control, we first demonstrated siRNA-based knockdown of *NFE2L2* mRNA – which would consequently reduce NMRAL2P expression – in A549 cells with published oligos (**112**) and saw efficient depletion of *NFE2L2* mRNA to ∼25% (30% in SKMES1 cells) (**Fig. 4F** and **Supplementary Fig. S11C**). Transfection of all three gapmers into A549 and SKMES1 cells resulted in knockdown of *NMRAL2P* expression to similar levels as *NFE2L2* knockdown (**Fig. 4G**, and **Supplementary Fig. S11D**). Notably, gapmer performance variability, although small, was proportional to average SHAPE reactivity of the targeted region. In-cell SHAPE probing thus enables more rapid (and less costly) identification of effective ASOs than iterative tiling, especially for long ncRNAs like NMRAL2P.

### Depletion of NMRAL2P inhibits NSCLC cell proliferation and sensitizes cells to cisplatin

Having established tools for NMRAL2P depletion and identified an association between higher NMRAL2P expression and poorer overall survival in lung adenocarcinoma, we used our gapmers to test whether NMRAL2P supports A549 (LUAD) and SKMES1 (LUSC) cell proliferation. In A549 cells, all three gapmers significantly reduced proliferation relative to the scrambled gapmer control, as did *NFE2L2* siRNA relative to the control siRNA (**Fig. 5A**). Because NRF2 activation can promote therapeutic resistance through induction of detoxification and drug efflux pathways (**113**), we next asked whether NMRAL2P depletion sensitizes A549 cells to cisplatin, a platinum-based chemotherapy used in first-line NSCLC treatment regimens. We generated cisplatin dose-response curves following treatment with NMRAL2P-targeting or scrambled gapmers (**Fig. 5B**) and *NFE2L2*-targeting or control siRNAs (**Fig. 5C**) and calculated the corresponding half-maximal inhibitory concentrations (IC_50_).

**Figure 5:**
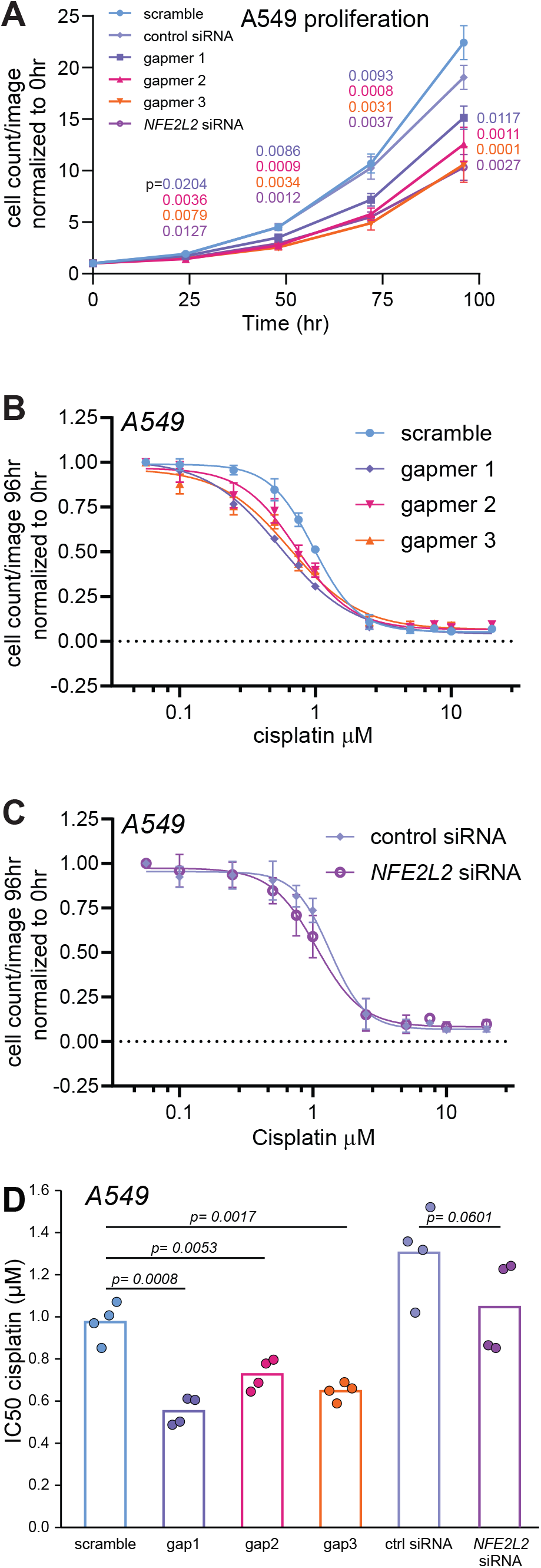
Depletion of NMRAL2P inhibits NSCLC cell proliferation and sensitizes cells to cisplatin. (**A**) Proliferation measurements of A549 cell counts over time with indicated oligo treatments. p-values for each time point compared to relevant controls are displayed. IC_50_ curves from A549 cells 96 hours post-transfection of indicated (**B**) NMRAL2P gapmer and scramble control or (**C**) *NFE2L2*-targeting and control siRNA treated with indicated cisplatin concentration. (**D**) IC_50_ values from B and C plotted with standard deviation and indicated significance. Error bars represent standard deviation. All p-values calculated from paired 2-sided t-tests. Proliferation and cisplatin proliferation assays were performed in 4 biological replicates.

NMRAL2P-targeting gapmers significantly lowered the cisplatin IC_50_ relative to the scrambled control. NFE2L2 siRNA also reduced the estimated IC_50_, although this change was not statistically significant relative to the control siRNA (**Fig. 5D**). Together with the proliferation assays, these results indicate that direct NMRAL2P depletion reduces A549 proliferation and increases cisplatin sensitivity under the conditions tested. We also tested the effects of NMRAL2P gapmer and *NFE2L2* siRNA on SKMES1 cells and saw similar trends to proliferation and cisplatin sensitivity (**Supplementary Fig. S12**). However, significant proliferation inhibition was more delayed in SKMES1 cells for both gapmers and *NFE2L2* siRNA, and the reductions in cisplatin IC_50_ were not significant.

### NMRAL2P depletion reveals a unique secondary transcriptional response

Because depletion of NMRAL2P ncRNA and *NFE2L2* mRNA produced similar effects on NSCLC proliferation, we used RNA-seq to assess whether these perturbations also elicited shared transcriptional responses (**Fig. 6**). Incomplete target depletion, particularly in SKMES1 cells, could reduce our ability to detect consistently differentially expressed genes using conventional significance thresholds. We therefore again used rank-rank hypergeometric overlap (RRHO) to identify concordant downregulation across conditions. We organized the comparisons into three groups: NMRAL2P depletion with three independent gapmers in A549 cells, NMRAL2P depletion with the same gapmers in SKMES1 cells, and NRF2 depletion with *NFE2L2* siRNA in both cell lines. Within each group, we used RRHO to compare each pair of treatment-versus-control gene rankings (**Supplementary Fig. S13**). We then retained genes shared by the regions of strongest concordant downregulation in all pairwise comparisons and intersected the resulting three group-level gene sets to identify genes concordantly downregulated across both NMRAL2P and NRF2 depletion conditions (**Fig. 6A**).

**Figure 6:**
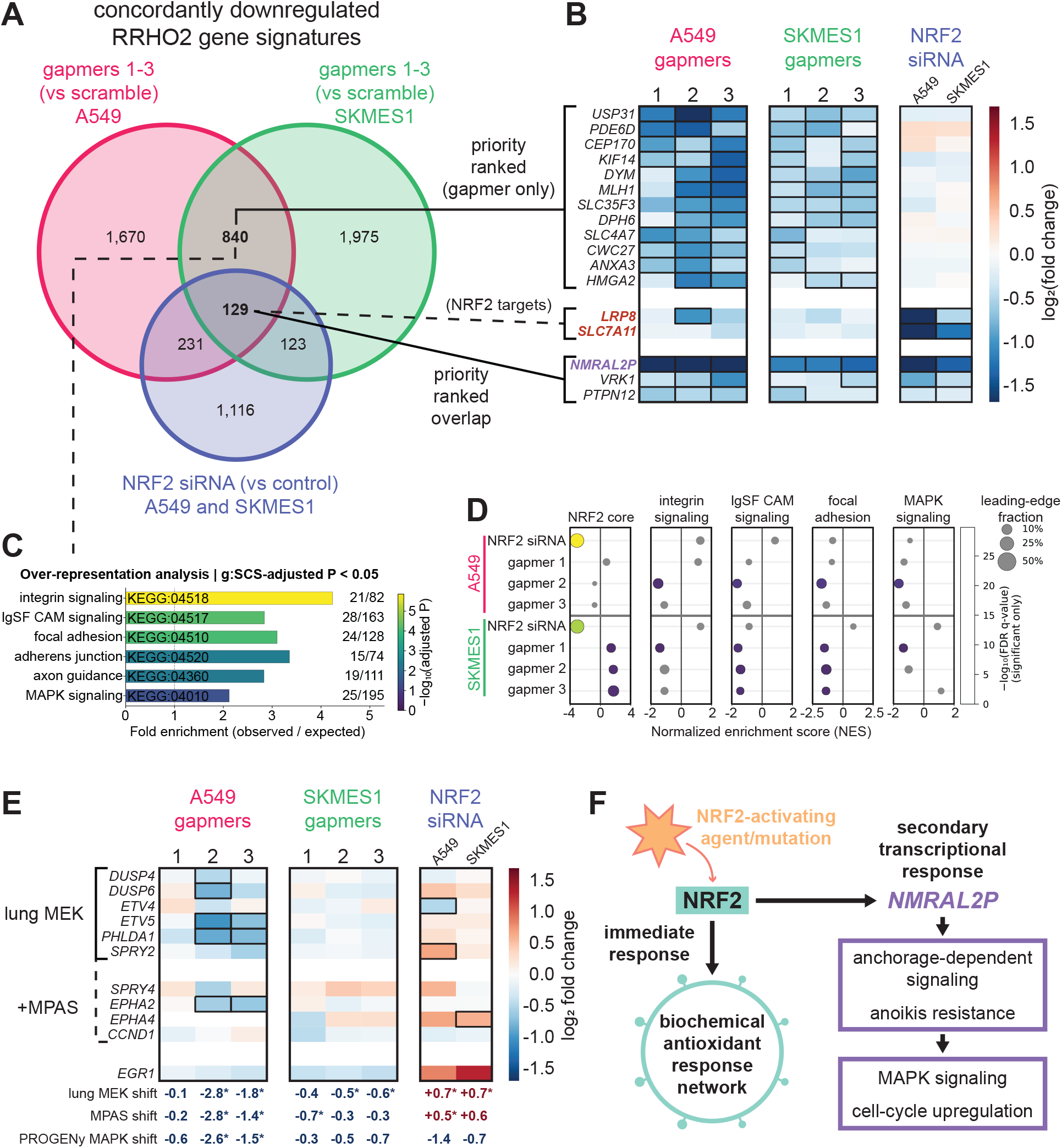
NMRAL2P depletion reveals a unique secondary transcriptional response. (**A**) Venn diagram representing intersection of indicated groups’ rank-rank hypergeometric overlaps. (**B**) Heat map of log_2_(fold change) for indicated genes comparing indicated gapmer- or siRNA-treated cells to corresponding control-treated cells. A border indicates a comparison achieved BH-adjusted P < 0.05. NRF2 core target gene names are colored red while noncoding RNA gene names are colored purple. Source of gene lists from Venn regions in A are indicated. With the exception of NRF2 targets, only genes significant in N-2 relevant samples are shown. (**C**) Enriched KEGG pathways from indicated Venn region. g:SCS = Set Counts and Sizes multiple testing correction from g:Profiler. Fractions of input genes overlapping total pathway genes measured are shown. (**D**) Gene set enrichment analysis for indicated pathways and treatments. (**E**) Heat maps as in B showing selected MAPK-associated genes. Changes in the indicated MAPK signature scores are expressed in standard deviation units relative to the corresponding control. * = BH-adjusted P < 0.05, from two-sided t-tests of treatment effects after accounting for differences between matched experimental replicates. (**F**) Model of NMRAL2P secondary transcriptional response.

Indeed, the number of significantly altered genes shared between all cell lines and gene depletions were few. In fact, the only gene significantly downregulated in every condition was NMRAL2P (**Fig. 6B**), confirming both our gapmers effectiveness and NRF2’s control of NMRAL2P expression. The only other two genes significantly downregulated in most conditions, VRK1 and PTPN12, encode regulators with potentially opposing effects on tumor progression. VRK1 supports cell-cycle progression and proliferation in lung cancer models (**114**), whereas PTPN12 can restrain oncogenic receptor tyrosine kinase signaling, including EGFR and HER2, and suppress tumor growth and metastasis in breast cancer models (**115**). There were a larger set of genes significantly downregulated exclusively in most NMRAL2P depleted samples: The highest-priority downregulated genes included USP31, which has been reported to support E2F1 stability and proliferation in lung squamous carcinoma cells (**116**); PDE6D, a positive regulator of KRAS localization and signaling (**117, 118**); and the proteins CEP170 and KIF14, involved in microtubule organization (**119**) and cytokinesis (**120**), respectively. Together with reduced expression of HMGA2 and ANXA3, established promoters of lung cancer proliferation (**121–123**), these changes identify plausible contributors to growth inhibition following NMRAL2P depletion.

Only two genes from the 42-gene NRF2 core set of targets (**73**), LRP8 and SLC7A11, were present in the shared downregulation signature across all three analysis groups (**Fig. 6B**). Four additional core genes were represented in the NMRAL2P-depletion signature in either A549 or SKMES1 cells, whereas the remaining 36 were restricted to the NRF2-depletion signature. This limited overlap argues against NMRAL2P broadly mediating the canonical NRF2 transcriptional response and instead supports a model in which NMRAL2P contributes to a distinct downstream gene-expression program.

To explore potential downstream expression programs without preselecting pathways, we performed over-representation analysis (ORA) against the full Kyoto Encyclopedia of Genes and Genomes (KEGG) pathway collection in g:Profiler (**84**), using genes shared between the A549 and SKMES1 NMRAL2P-depletion signatures but absent from the NRF2-depletion signature (**Fig. 6C**). The most significantly enriched pathways included integrin signaling, IgSF CAM signaling, focal adhesion, adherens junction, axon guidance, and MAPK signaling, suggesting that NMRAL2P depletion preferentially affects genes involved in cell adhesion, cytoskeletal organization, and growth-factor signaling. To determine whether these pathway signatures differed between NMRAL2P and NRF2 depletion, we performed targeted gene set enrichment analysis using the complete ranked gene lists from each RNA-seq comparison (**Fig. 6D**). The NRF2 core gene set showed selective negative enrichment following NRF2 depletion, whereas the pathways identified by over-representation analysis showed more pronounced negative enrichment following NMRAL2P depletion. These findings further support a model in which NMRAL2P contributes to a distinct transcriptional program downstream of NRF2.

To test whether reduced MAPK output could contribute to the proliferation defect following NMRAL2P depletion, we evaluated three complementary transcriptional readouts: a lung MEK inhibitor six-gene signature (**85**), a ten-gene MAPK Pathway Activity Score (MPAS) (**120**), and the weighted PROGENy MAPK signature (**120**) (**Fig. 6E** and **Supplementary Fig. S14**). All three scores decreased significantly in A549 cells treated with gapmers 2 and 3, whereas responses in gapmer-treated SKMES1 cells reached significance for selected lung MEK and MPAS comparisons. In contrast, NRF2 depletion did not significantly reduce any MAPK score in either cell line and significantly increased the lung MEK and MPAS scores in some cases. Across the broader PROGENy pathway panel, NMRAL2P depletion consistently reduced EGFR- and MAPK-associated transcriptional output and increased JAK-STAT-associated output in both cell lines, with other responses varying by cell line (**Supplementary Fig. S15**). Consistent with reduced MAPK-associated transcriptional output, the immediate-early response gene *EGR1* showed concordant, although not statistically significant, downregulation across NMRAL2P gapmer comparisons in both cell lines. Together, these results support reduced MAPK transcriptional output as a potential contributor to NMRAL2P-dependent growth inhibition and indicate a selective signaling response distinct from that elicited by NRF2 depletion.

Together, these findings support a working model in which NRF2 activates both a canonical antioxidant defense program and an NMRAL2P-dependent secondary response that supports tumor-cell growth (**Fig. 6F**). The limited overlap between the NMRAL2P-depletion signatures and the NRF2 core gene set suggests that NMRAL2P preferentially sustains a distinct transcriptional program. Enrichment of adhesion- and cytoskeleton-associated pathways by ORA (**Fig. 6C**), their negative enrichment following NMRAL2P depletion by GSEA (**Fig. 6D**), and reduced EGFR/MAPK-associated output inferred by PROGENy (**Supplementary Fig. S15**) collectively implicate this program in coupling cell matrix interactions to proliferative signaling. Downregulation of genes involved in mitogenic signaling and cell division further supports a role in maintaining cell-cycle progression. We therefore propose that NMRAL2P helps sustain signaling that enables anchorage-independent growth and resistance to anoikis, complementing NRF2’s broader biochemical antioxidant response. The specific contributions of altered MAPK signaling and anoikis resistance remain mechanistic hypotheses requiring direct functional validation.

## DISCUSSION

This study identifies NMRAL2P as an NRF2-induced, predominantly nuclear long noncoding RNA expressed as three major isoforms in NSCLC (**Fig. 2** and **Fig. 3**). NMRAL2P depletion inhibited proliferation and growth-associated transcriptional programs without broadly reproducing the loss of canonical NRF2 antioxidant gene expression (**Fig. 6**), supporting its role in a selective downstream response to NRF2 activation. Notably, NMRAL2P was not appreciably expressed in NRF2-inactive cells among the models examined. This expression pattern raises the possibility of preferentially targeting NRF2-active cancer cells while sparing NRF2-inactive cells, although selectivity would need to be established in normal tissues, including those undergoing physiological NRF2 activation. NMRAL2P depletion inhibited proliferation and increased cisplatin sensitivity in NSCLC cells (**Figs. 5** and **Supplementary Fig. S12**). In TCGA-LUAD, higher NMRAL2P expression was associated with poorer overall survival within NRF2-active patient cohorts (**Fig. 2**), supporting its potential clinical relevance. Together with its limited expression in NRF2-inactive cells, these findings motivate further investigation of NMRAL2P as a therapeutic target in NRF2-active lung adenocarcinoma. In-cell chemical probing also provided a practical means of identifying sites for effective gapmer-mediated depletion of this nuclear RNA (**Fig. 4**).

### Coding and regulatory divergence may have contributed to the evolution of NMRAL2P as a nuclear lncRNA

Comparative sequence analysis identifies in *NMRAL2P* a conserved NMRAL1-related ORF interrupted by a human-specific premature stop, suggesting a recent change in NMRAL2P coding potential (**Fig. 3**). The protein-coding ancestry of XIST and other X-inactivation-associated lncRNAs provides precedent for the emergence of regulatory noncoding RNAs through loss of ancestral coding potential (**124, 125**). Regulatory divergence may also have contributed to NMRAL2P function: while human NMRAL2P is strongly NRF2-responsive in NSCLC cells, neither human *NMRAL1* (this study) nor mouse *Nmral2p* (**106**) are NRF2-responsive at the RNA level. Expression of *NMRAL2P* has not been demonstrated in other primates, although NRF2-active models were not tested. These observations suggest that acquisition of NRF2-responsive regulation may have accompanied NMRAL2P’s divergence toward a functional nuclear RNA. RNA surveillance provides a possible connection between coding disruption and localization. Endogenous transcripts containing detained introns frequently harbor premature termination codons yet remain nuclear and escape NMD, demonstrating that nuclear detention can preserve potentially NMD-sensitive RNA (**126**). However, these studies implicate incomplete splicing rather than stop codon recognition as the basis for detention and therefore do not establish how the premature stop in spliced NMRAL2P affects its localization. The evolutionary order of regulatory change, coding disruption, and nuclear localization remains unresolved; direct investigation is needed to determine whether the human-specific stop affects translation, NMD susceptibility, or subcellular distribution.

### NMRAL2P supports context-dependent regulatory programs beyond canonical NRF2 antioxidant responses

Although NRF2 is best characterized as a regulator of antioxidant and detoxification enzymes, sustained NRF2 activation also influences broader cellular programs through downstream regulatory factors, whose contributions to cell survival and proliferation remain less well defined. NRF2 directly activates *NOTCH1* transcription, and restoration of Notch signaling rescues impaired liver regeneration in NRF2-deficient mice (**127**), linking NRF2 to proliferative tissue repair. Separately, NRF2-dependent induction of the transcriptional regulator ATF3 suppresses IL-6 production in endotoxin-stimulated monocytes (**128**), demonstrating regulation of inflammatory responses through a secondary transcriptional effector. In our datasets, *NOTCH1* and *ATF3* expression also decreased following NRF2 depletion in both A549 and SKMES1 cells, although these changes did not reach statistical significance. These examples provide precedent for a model in which NRF2-induced NMRAL2P mediates a secondary regulatory program supporting tumor cell growth. Decreases in established, perturbation-validated MAPK activity signatures following NMRAL2P depletion (**Figs. 6** and **Supplementary S14–S15**) support a role for NMRAL2P in sustaining MAPK signaling, although ERK kinase activity was not directly measured and the underlying mechanism remains unresolved.

The selective effects of NMRAL2P depletion are consistent with previous reports connecting NMRAL2P/LOC344887 to growth-associated phenotypes. In gallbladder cancer, LOC344887 depletion reduced proliferation, migration, invasion, and mesenchymal marker expression (**129**). In hepatocellular carcinoma, LOC344887 sustained STAT3 signaling and promoted HMGA2 expression (**130**). NMRAL2P depletion likewise reduced HMGA2 expression in our datasets, but increased PROGENy JAK-STAT activity scores (**Figs. 6** and **Supplementary Fig. S15**). Thus, the direction of HMGA2 regulation agrees with the previous report, whereas the STAT-associated response differs. Although the underlying mechanisms may differ across cellular contexts, these findings collectively support a contribution of NMRAL2P to tumor-associated gene regulation. By contrast, despite its reported role in NQO1 induction in sulforaphane-treated colon cancer cells (**50**) and GPX2 transcription through association with NRF2 in head-and-neck cancer cells (**52**), our results do not support a broad requirement for NMRAL2P in maintaining the core NRF2 antioxidant transcriptional program in NSCLC. Consistent with this distinction, NMRAL2P depletion in lung fibroblasts impaired the antifibrotic response to sulforaphane without affecting NQO1 induction (**51**). NMRAL2P may therefore function as a selective downstream effector of NRF2, supporting growth-associated programs without broadly mediating its canonical antioxidant response.

### Distinct survival associations in LUAD and LUSC do not exclude therapeutic potential in either histology

The weaker responses to NMRAL2P depletion in SKMES1 than in A549 further suggest that its contribution may depend on cellular context. SKMES1 showed a smaller reduction in NMRAL2P upon gapmer knockdown (**Supplementary Fig. S11**), more delayed inhibition of proliferation, and no statistically significant change in cisplatin IC_50_ under the conditions tested (**Supplementary Fig. S12**). Poorer transfection efficiency or lower gapmer activity in SKMES1 could contribute to the reduced knockdown and consequently weaker phenotypic responses. Dose- and time-response studies that achieve comparable NMRAL2P depletion would help determine whether these differences reflect knockdown efficiency or distinct biological requirements for NMRAL2P.

The patient data likewise support evaluating NMRAL2P separately in LUAD and LUSC. Higher NMRAL2P expression was associated with poorer survival in NRF2-active LUAD cohorts, whereas comparable associations were not detected in LUSC (**Figs. 2** and **Supplementary Fig. S5**). However, the absence of a survival association does not exclude a therapeutic benefit from NMRAL2P depletion in LUSC: DepMap collections (26Q1) identify similar NRF2 dependence in both LUAD (20/53 lines, 37.7%) and LUSC (12/21 lines, 57.1%), including both A549 and SKMES1. Moreover, NMRAL2P expression was significantly higher across the TCGA LUSC cohort than in LUAD (**Supplementary Fig. S5G**), and its location at 3q27.2 falls within the recurrently amplified 3q region in lung squamous cancers (**131**). Copy-number gain could contribute to elevated expression, although that relationship requires direct assessment. Together with the less efficient depletion achieved in SKMES1, these findings support further investigation of NMRAL2P targeting in LUSC. Comparisons using matched depletion across additional models would help establish whether sensitivity varies with histology, NRF2 activity, NMRAL2P abundance, or other molecular features.

### In-cell RNA accessibility measurements can guide effective gapmer selection and inform predictive ASO design

Finally, the accessibility-guided gapmer workflow offers a practical route for testing nuclear lncRNA function and prioritizing therapeutic candidates. In-cell SHAPE-MaP identified locally flexible regions shared among NMRAL2P isoforms, enabling selection of gapmers that achieved approximately 80-90% depletion (**Fig. 4, Supplementary Figs. S10 and S11**). This approach complements computational prediction (**132**) and oligonucleotide tiling (**133**) by providing experimental information from RNA in its cellular environment. Structural accessibility has long been recognized as a determinant of antisense activity, and recent work has directly correlated SHAPE-derived accessibility with gapmer efficacy in viral RNA (**134**). Our findings extend the utility of this strategy to a predominantly nuclear human lncRNA. A larger dataset pairing in-cell reactivity measurements with ASO dose-response profiles could establish whether accessibility across the complete target site predicts knockdown potency and maximal depletion. Combining these measurements with sequence, hybridization, and oligonucleotide-chemistry features could support models that prioritize effective candidates before extensive screening. Validation across independent transcripts and cell types would determine how broadly such models transfer. By reducing the number of candidates requiring experimental optimization, this strategy could accelerate functional interrogation of nuclear lncRNAs and improve the efficiency of early ASO therapeutic testing.

## ACKNOWLEDGEMENTS

M. Ben Major lab for the H1299-YFP-NQO1 cells and pHAGE-UB-FLAG-GFP plasmid. Kira E. Holton for her cellular fractionation protocol.

## AUTHOR CONTRIBUTIONS

Jae A. Bucknor: Conceptualization, Data curation, Formal analysis, Funding acquisition, Investigation, Methodology, Validation, Visualization, Writing – original draft, Writing – review & editing. Brittany M. Bowman: Conceptualization, Data curation, Formal analysis, Investigation, Methodology, Supervision, Validation, Visualization, Writing – original draft, Writing – review & editing. Shannon Stein: Conceptualization, Methodology, Writing – review & editing. Chase A. Weidmann: Conceptualization, Data curation, Formal analysis, Funding acquisition, Methodology, Project administration, Resources, Software, Supervision, Visualization, Writing – original draft, Writing – review & editing.

## SUPPLEMENTARY DATA

Supplementary Data are available online.

## CONFLICT OF INTEREST

Authors declare no competing interests.

## FUNDING

This work was supported by the National Institutes of Health [CA262349 and GM155542 to C.A.W, GM245470 to J.A.B]; and by the University of Michigan [Rackham Graduate Student Research Grant to J.A.B. and S.S., Rackham Graduate School PACE Fellowship To J.A.B and C.A.W.]. Funding for open access charge: National Institutes of Health.

## DATA AVAILABILITY

The raw sequencing reads and processed data for the ribosomal RNA-depleted bulk RNA-seq, Plasmidsaurus 3′ RNA-seq, and SHAPE-MaP experiments can be found in the NCBI Gene Expression Omnibus (GEO) under GSE354488, GSE354489, and GSE354490.

## Notes

### Competing Interest Statement

The authors have declared no competing interest.

