## Supplementary_Figures for "Accessibility-guided targeting of NRF2-responsive long noncoding RNA NMRAL2P alters a distinct gene expression program and reduces non-small cell lung cancer proliferation"

#### Contents

Supplementary figures and legends

### Figure S1

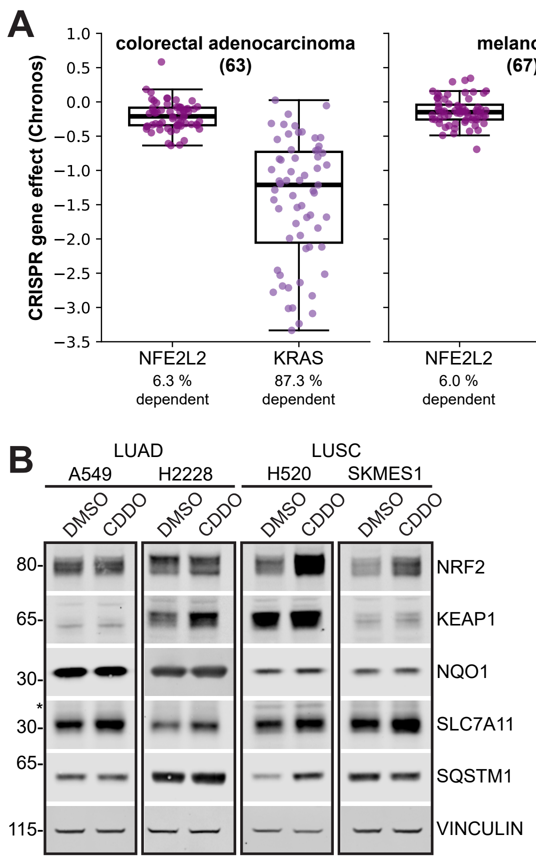
A
colorectal adenocarcinoma (63)
melanoma (67)
invasive breast carcinoma (50)
pancreatic adenocarcinoma (46)
CRISPR gene effect (Chronos)
1.0
0.5
0.0
−0.5
−1.0
−1.5
−2.0
−2.5
−3.0
−3.5
NFE2L2 6.3 % dependent
KRAS 87.3 % dependent
NFE2L2 6.0 % dependent
BRAF 74.6 % dependent
NFE2L2 6.0 % dependent
ERBB2 36.0 % dependent
NFE2L2 15.2 % dependent
KRAS 95.7 % dependent
B
LUAD
A549
H2228
LUSC
H520
SKMES1
DMSO
CDDO
80-
65-
30-
*
30-
65-
115-
NRF2
KEAP1
NQO1
SLC7A11
SQSTM1
VINCULIN

**Figure S1:** Additional *NFE2L2* cell line dependencies and responses to CDDO-Me. (A) Cancer Dependency Map (26Q1) gene effects for indicated cell types from knockout of *NFE2L2* or indicated cancer type-relevant gene. Boxes show the median and interquartile range; whiskers extend to the most extreme observations within 1.5× the interquartile range. (B) Western blots showing NRF2, KEAP1 and NRF2 target protein levels comparing lung adenocarcinoma (LUAD) and squamous cell carcinoma (LUSC) cells to cells treated with CDDO-Me. Vinculin is a loading control.

### Figure S2

#### RRHO2 pairwise comparison heatmaps

##### Cancer

A549 vs HBE vs H2228 vs HBE

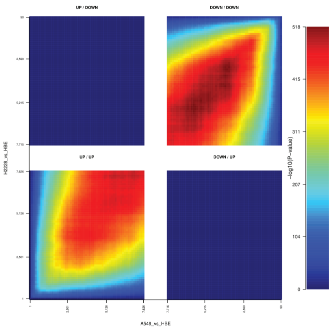

A549 vs HBE vs H520 vs HBE

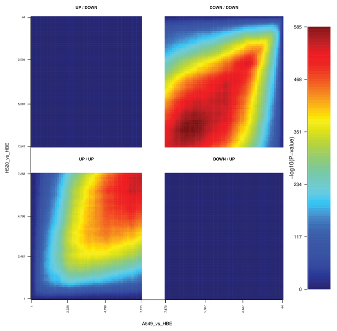

A549 vs HBE vs SKMES1 vs HBE

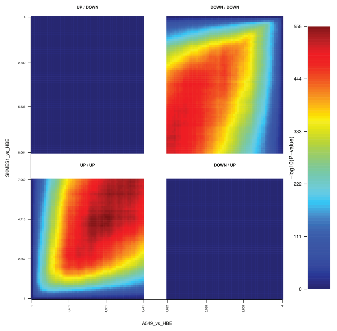

H2228 vs HBE vs H520 vs HBE

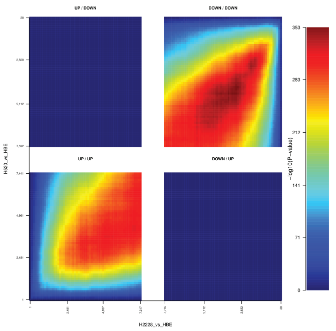

H2228 vs HBE vs SKMES1 vs HBE

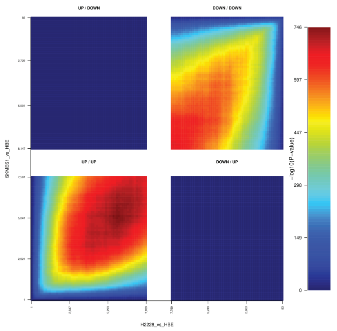

H520 vs HBE vs SKMES1 vs HBE

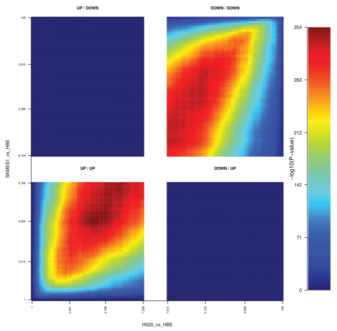

##### Treatment

CDDO vs CALU1 vs CDDO vs H1299

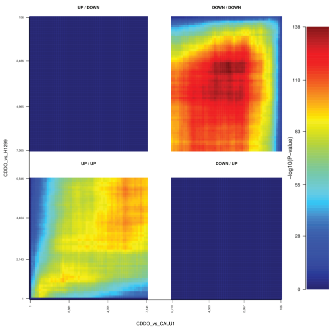

CDDO vs CALU1 vs CDDO vs HBE

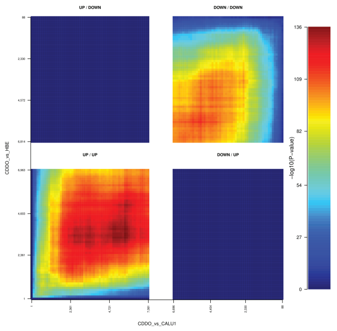

CDDO vs H1299 vs CDDO vs HBE

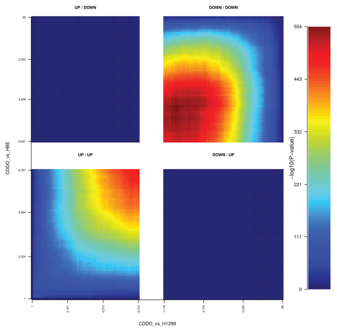

##### Mutation

E79Q vs H1299 vs KEAP1KO vs H1299

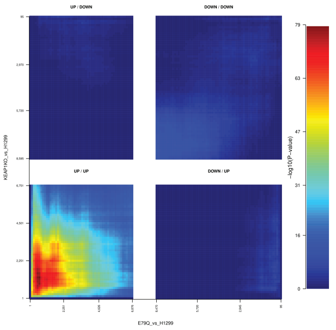

**Figure S2. Pairwise overlap of ranked differential gene expression profiles in lung cell models of NRF2 activation.** Rank–rank hypergeometric overlap (RRHO2) heatmaps compare differential expression profiles across three sets of contrasts. **Cancer:** all six pairwise comparisons among A549, H2228, H520, and SKMES1 cells, each profiled relative to human bronchial epithelial (HBE) cells. **Treatment:** all three pairwise comparisons of CDDO treatment responses in CALU-1, H1299, and HBE cells. **Mutation:** comparison of the H1299-NRF2$^{E79Q}$ and H1299-KEAP1$^{sgRNA}$ (knockout) expression profiles, each relative to H1299 control cells. The axes represent rank cutoffs in the two differential expression profiles. The lower-left and upper-right quadrants show overlap among concordantly upregulated and concordantly downregulated genes, respectively. The upper-left and lower-right quadrants show overlap among genes regulated in opposite directions. Colors indicate $-\log_{10}$ of nominal hypergeometric overlap P values, with a separate color scale for each heatmap.

### Figure S3

#### NRF2 core gene set | GSEA enrichment

##### Cancer

###### A549 vs HBE

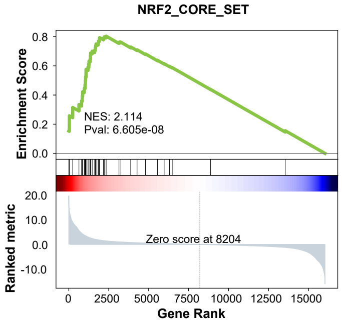


###### SKMES1 vs HBE

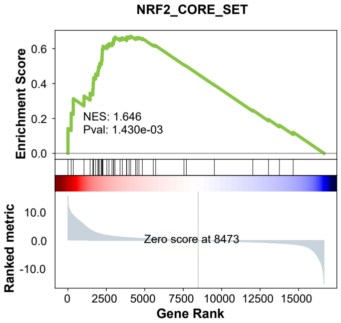


###### H2228 vs HBE

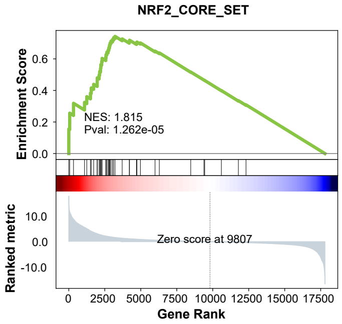


###### H520 vs HBE

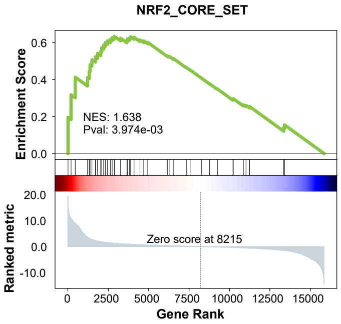


##### Treatment

###### CDDO vs HBE

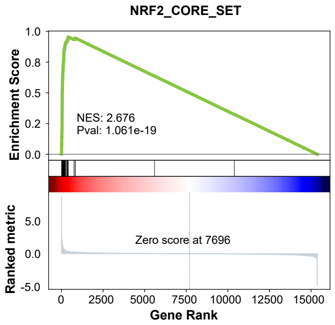


###### CDDO vs H1299

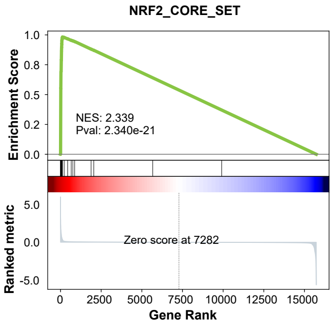


###### CDDO vs CALU1

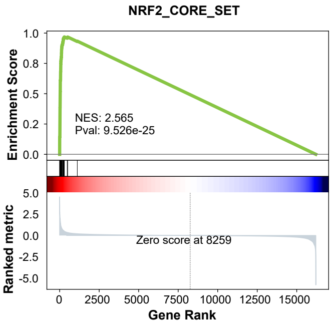


##### Mutation

#### E79Q vs H1299

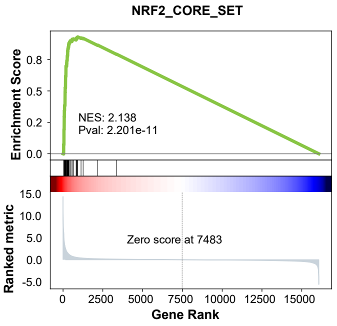


###### KEAP1KO vs H1299

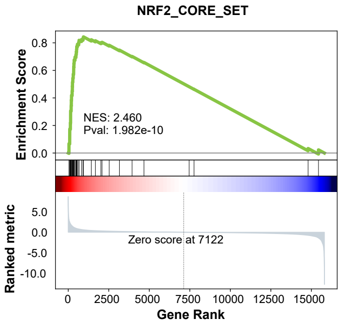

**Figure S3. The NRF2 core gene set is positively enriched across lung cancer, CDDO treatment, and genetic perturbation models.** Gene set enrichment analysis (GSEA) was performed on ranked differential expression profiles using a 42-gene NRF2 core target gene set. The plots show A549, SKMES1, H2228, and H520 cells each compared with human bronchial epithelial (HBE) cells (Cancer), CDDO-treated HBE, H1299, and CALU-1 cells each compared with their respective controls (Treatment), and H1299-NRF2$^{E79Q}$ and H1299-KEAP1$^{sgRNA}$ (knockout) cells each compared with H1299 control cells (Mutation). Green curves show the running enrichment score, black ticks mark the positions of NRF2 core genes in each ranked list, and gray profiles show the ranking metric. The dashed vertical line marks the rank at which the metric crosses zero. Normalized enrichment scores (NES) and nominal *P* values are shown for all contrasts.

### Figure S4

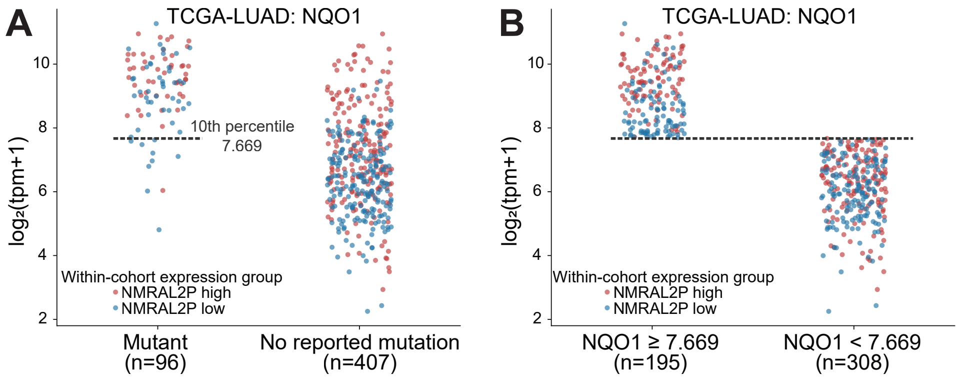

A
TCGA-LUAD: NQO1
log₂(tpm+1)
10th percentile
7.669
Within-cohort expression group
NMRAL2P high
NMRAL2P low
Mutant
(n=96)
No reported mutation
(n=407)
B
TCGA-LUAD: NQO1
log₂(tpm+1)
Within-cohort expression group
NMRAL2P high
NMRAL2P low
NQO1 ≥ 7.669
(n=195)
NQO1 < 7.669
(n=308)

**Figure S4. NQO1 expression in TCGA lung cancer cohorts. (A)** NQO1 expression in lung adenocarcinoma (TCGA-LUAD) tumors grouped by the presence (left) or absence (right) of a reported activating mutation in *NFE2L2* or *KEAP1*. Dashed lines mark the 10th percentile of NQO1 expression in the mutation-positive group. **(B)** The same tumors grouped by whether NQO1 expression was at or above the corresponding cutoff (7.669). Each point represents one tumor. Red and blue indicate top and bottom 50 percent of tumors by NMRAL2P expression, respectively, based on the median within each subgroup. Number of included patient tumors in each group is indicated.

### Figure S5

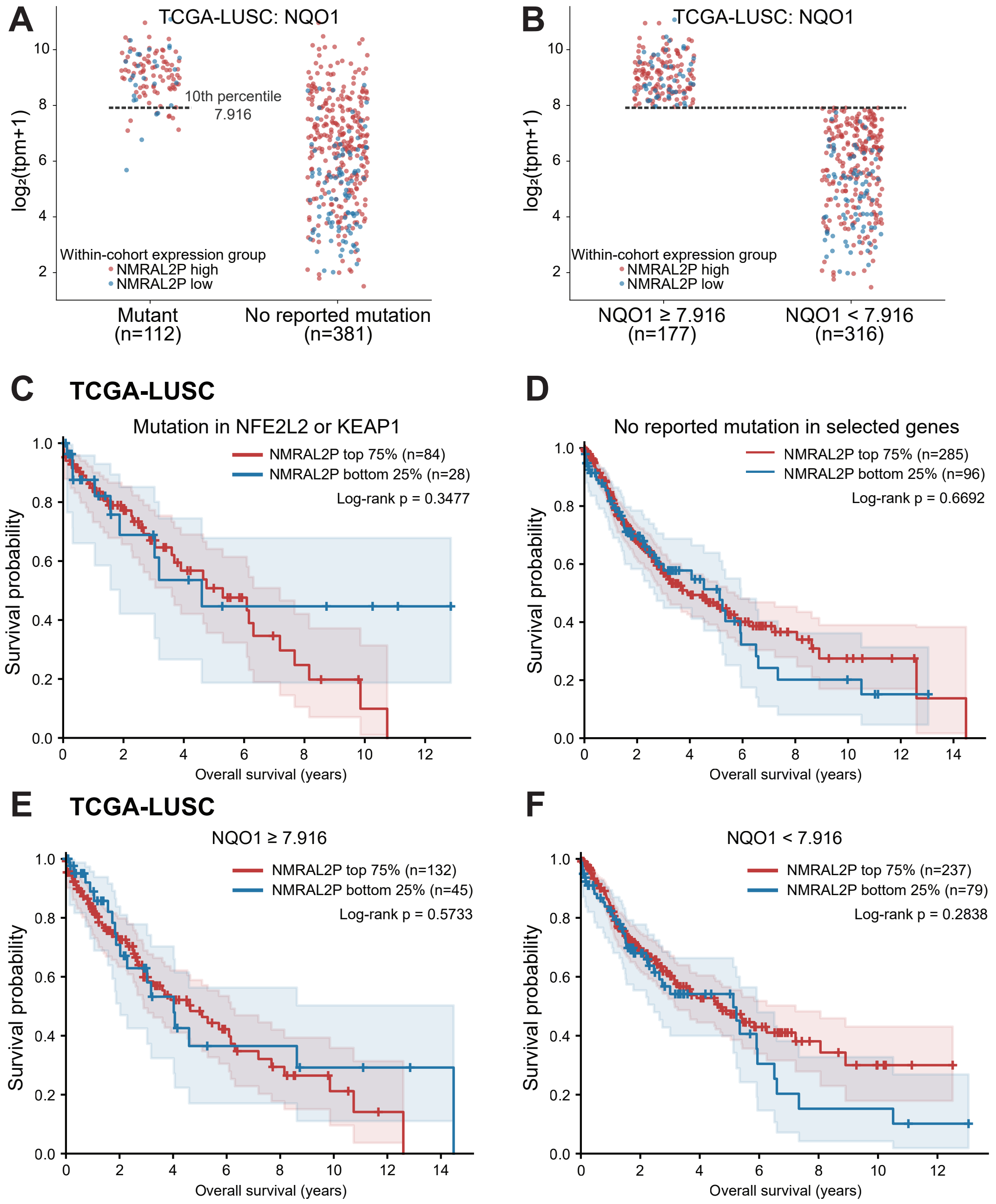


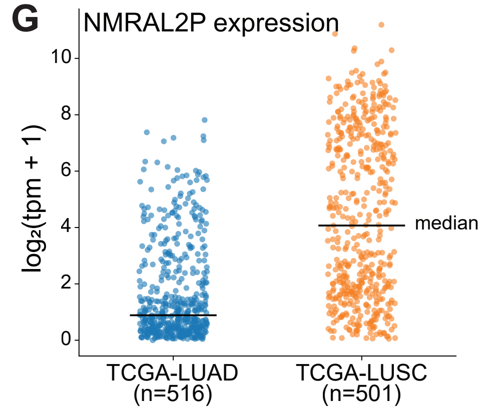


**H** **Complete-case Cox proportional hazards models**

| Cancer | Variables included in model | NMRAL2P HR (95% CI) | P |
|---|---|---|---|
| LUAD | NMRAL2P | 1.20 (1.05–1.39) | 0.0092 |
| LUAD | NMRAL2P + mutation category | 1.25 (1.04–1.50) | 0.0148 |
| LUAD | NMRAL2P + NQO1 category | 1.31 (1.09–1.59) | 0.0046 |
| LUAD | NMRAL2P + age + sex + stage | 1.16 (1.00–1.35) | 0.0499 |
| LUAD | NMRAL2P + mutation category + age + sex + stage | 1.18 (0.98–1.43) | 0.0849 |
| LUAD | NMRAL2P + NQO1 category + age + sex + stage | 1.26 (1.03–1.54) | 0.0242 |
| LUSC | NMRAL2P | 0.93 (0.81–1.06) | 0.2678 |
| LUSC | NMRAL2P + mutation category | 0.94 (0.80–1.10) | 0.4225 |
| LUSC | NMRAL2P + NQO1 category | 0.90 (0.76–1.08) | 0.2571 |
| LUSC | NMRAL2P + age + sex + stage | 0.95 (0.83–1.09) | 0.4337 |
| LUSC | NMRAL2P + mutation category + age + sex + stage | 0.96 (0.82–1.12) | 0.5844 |
| LUSC | NMRAL2P + NQO1 category + age + sex + stage | 0.91 (0.76–1.08) | 0.2857 |

**Figure S5. LUSC cohort NQO1 expression and survival, LUAD and LUSC NRMAL2P expression, and Cox regression modeling. (A)** NQO1 expression in TCGA lung squamous cell carcinoma (LUSC) tumors grouped by the presence of a reported activating mutation in *NFE2L2* or *KEAP1*. The dashed line marks the 10th percentile of NQO1 expression in the mutation-positive group (7.916). **(B)** The same tumors grouped by NQO1 expression at or above versus below 7.916. In A and B, each point represents one tumor, and red and blue indicate high (top 75%) and low (bottom 25%) NMRAL2P expression within the indicated subgroup. Kaplan–Meier estimates of overall survival by NMRAL2P expression in TCGA-LUSC patients **(C)** with a reported *NFE2L2* or *KEAP1* mutation, **(D)** without a reported mutation in either gene, **(E)** with NQO1 expression ≥7.916, or **(F)** with NQO1 expression <7.916. Shaded regions indicate pointwise 95% confidence intervals, tick marks indicate censored observations, and *P* values are from log-rank tests. **(G)** NMRAL2P expression in TCGA lung adenocarcinoma (LUAD) and LUSC tumors, shown as $\log_2$ (TPM + 1). Horizontal lines indicate cohort medians. **(H)** Complete-case Cox proportional hazards models for overall survival in LUAD and LUSC. The table reports hazard ratios for NMRAL2P expression, 95% confidence intervals, and *P* values for the indicated covariate sets.

### Figure S6

```
                  1                                                50
human             MASKKVITVFGATGAQGGSVARAILESKKFAVRAVTRDVT*PNALELQRL
chimp             ........................................R.........
gorilla           ........................................R.........
rhesus macaque    ........................................R.K.......
gray mouse lemur  ..N....A..................N.......L.....G.K..A.RD.

                  51                                              100
human             GAEVVKGDLNDKASVDSALKGVYGAFLVTNFWDPLNQDKEVCRGKLVADS
chimp             .............L............................Q.......
gorilla           ..........................................Q.......
rhesus macaque    .....................A..V........A..K....FQ......T
gray mouse lemur  ...........ET..EA................D.SIK....Q..M..QV

                  101                                             150
human             AKHLGLKHVVYSGLENVKRLTDGKLEVPHFDSKGEVEEYFWSIGIPMTSV
chimp             ............................................V.....
gorilla           ..................................................
rhesus macaque    .........................Q....................T...
gray mouse lemur  ..R..................G.........G.........AT.V.....

                  151                                             200
human             RVAAYFENFLAAWRPVKASDGDYYTLAVPMGDVPMDGISVADIGAAVSSI
chimp             ..................................................
gorilla           ...........................I...................C..
rhesus macaque    ..........T.....................................N.
gray mouse lemur  ..........T..K.A..P.......VL.................V....

                  201                                             250
human             FNSPEEFLGKAVGLSAEALTIQQYADVLSKVLGKEVRDAKITPEAFEKLG
chimp             .............................N....................
gorilla           ..................................................
rhesus macaque    ....................V.............K.........V.....
gray mouse lemur  .......I.........K............C...DI.........YK...

                  251                                             300
human             FPAAKEIANMCRFYEMKPDRDVNLTHQLNPKVKSFSQFISENQGAFKGM*
chimp             ......................K...........................
gorilla           ......................K...........................
rhesus macaque    ........D.....Q.......K...R.......................
gray mouse lemur  ..V.E.V....L..QT......K...R...............KE...D..
```

**Figure S6. A premature stop codon interrupts the human NMRAL2P reading frame.** Predicted amino acid sequences from the aligned NMRAL2P reading frame are shown for human, chimpanzee, gorilla, rhesus macaque, and gray mouse lemur. The human sequence is shown in full. Dots indicate matches to the human reference, letters indicate amino acid differences, and asterisks denote stop codons. At position 41, the human sequence contains a stop codon, whereas the corresponding codon encodes arginine in chimpanzee, gorilla, and rhesus macaque and glycine in gray mouse lemur. Purple underlining marks two putative human open reading frames, one beginning at the first methionine and ending at the premature stop and the other beginning at the downstream methionine at position 147 and ending at the stop at position 300.

### Figure S7

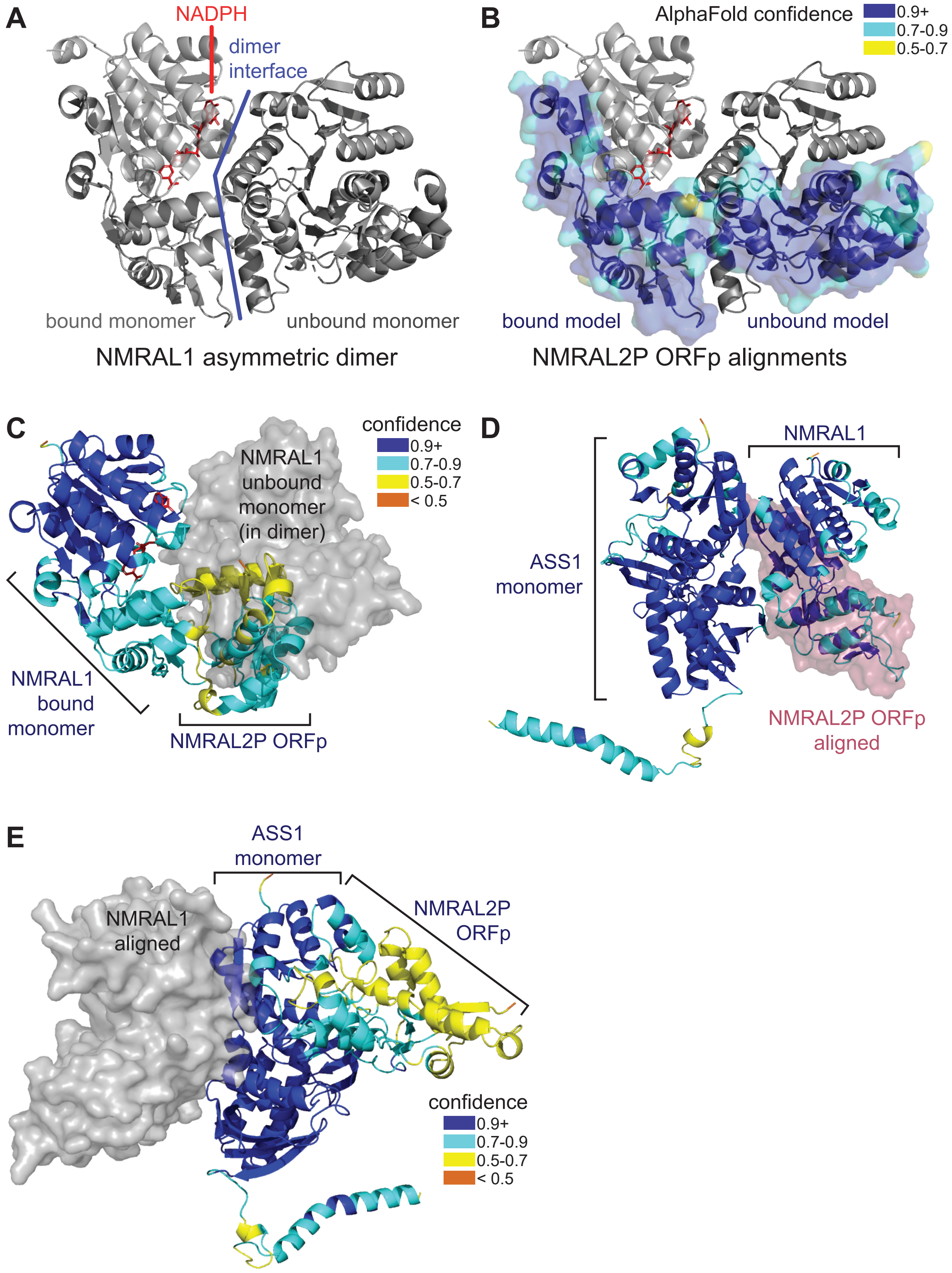
A
NADPH
dimer interface
bound monomer
unbound monomer
NMRAL1 asymmetric dimer
B
AlphaFold confidence
0.9+
0.7-0.9
0.5-0.7
bound model
unbound model
NMRAL2P ORFp alignments
C
NMRAL1 unbound monomer (in dimer)
confidence
0.9+
0.7-0.9
0.5-0.7
< 0.5
NMRAL1 bound monomer
NMRAL2P ORFp
D
NMRAL1
ASS1 monomer
NMRAL2P ORFp aligned
E
ASS1 monomer
NMRAL1 aligned
NMRAL2P ORFp
confidence
0.9+
0.7-0.9
0.5-0.7
< 0.5

**Figure S7. Limited evidence for NMRAL1-like function of major NMRAL2P open reading frame product (ORFp). (A)** Structure of NMRAL1 asymmetric dimer (PDB: 2EXX), with NADPH (red) pocket and dimerization interface (blue) indicated. **(B)** NMRAL1 structure as in A with surface projections of AlphaFold3 models of the NMRAL2P ORFp aligned to the bound and unbound NMRAL1 molecules. Surface model is colored by AlphaFold confidence scores. Alphafold multimer models of **(C)** NMRAL1 bound to NMRAL2P ORFp, **(D)** NMRAL1 bound to ASS1, and **(E)** NMRAL2P ORFp bound to ASS1 are shown colored by confidence score. Relative positions of NMRAL1 (gray) or NMRAL2P ORFp (red) aligned to each structure are represented as surface projections.

### Figure S8

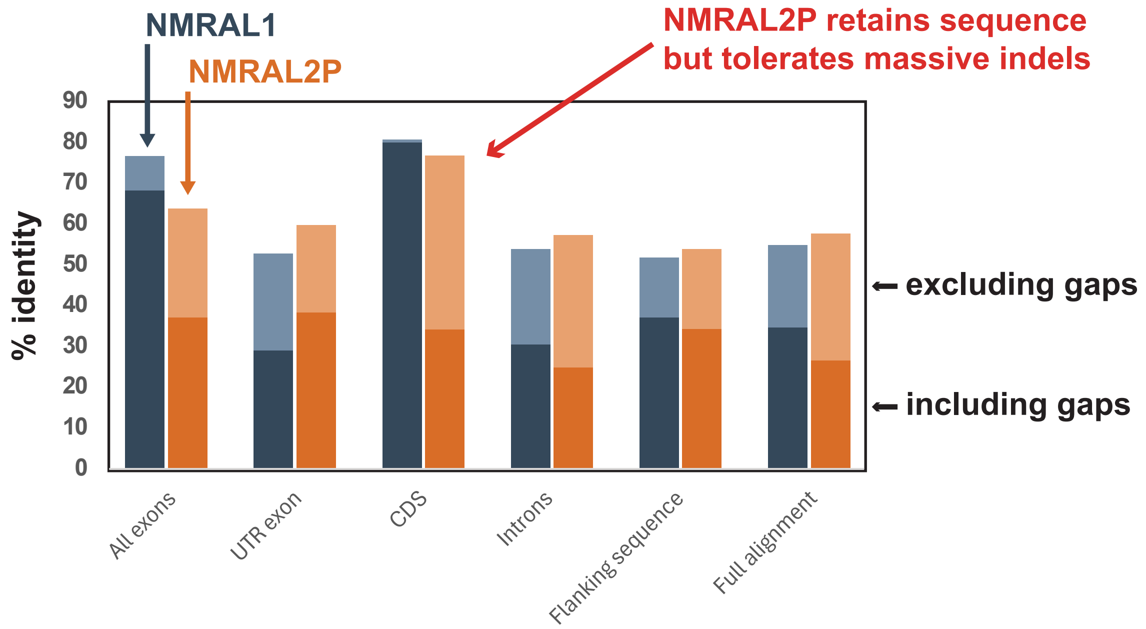

NMRAL1
NMRAL2P
NMRAL2P retains sequence but tolerates massive indels
% identity
90
80
70
60
50
40
30
20
10
0
excluding gaps
including gaps
All exons
UTR exon
CDS
Introns
Flanking sequence
Full alignment

**Figure S8. Greater divergence in the ancestral coding region of NMRAL2P supports its evolution into a noncoding RNA.** Percent nucleotide identity was calculated from pairwise human-mouse alignments of the NMRAL1 (blue) and NMRAL2P (orange) loci across the indicated regions. Dark bar segments show identity with gap positions included in the denominator, and the full bar heights show identity with gaps excluded. The region corresponding to the ancestral coding sequence shows similar gap-excluded identity at both loci, but including gaps reduces identity much more at NMRAL2P than at NMRAL1, consistent with relaxed constraint on the ancestral coding sequence.

### Figure S9

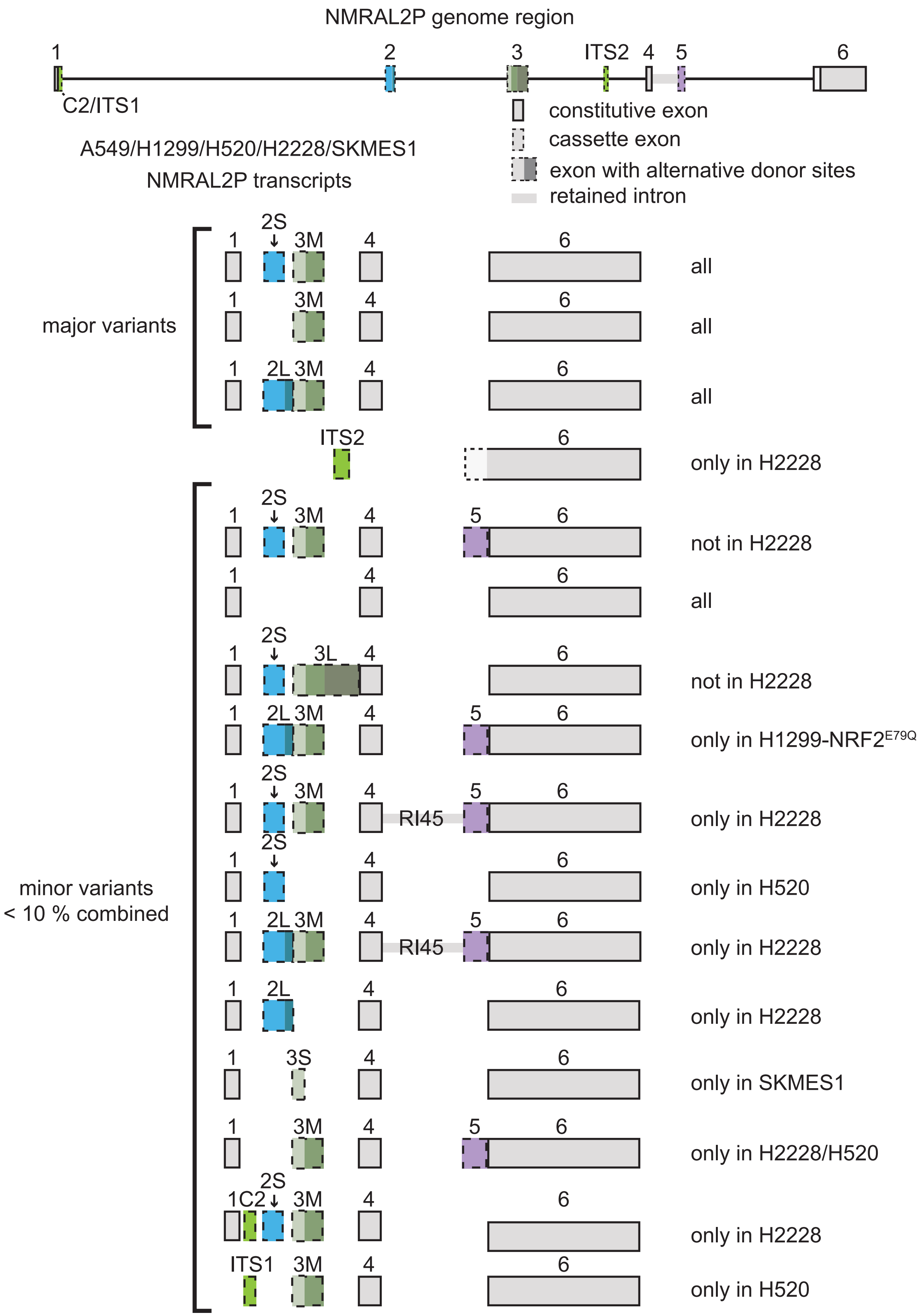
NMRAL2P genome region
1
2
3
ITS2
4
5
6
C2/ITS1
constitutive exon
cassette exon
exon with alternative donor sites
retained intron
A549/H1299/H520/H2228/SKMES1
NMRAL2P transcripts
major variants
minor variants
< 10 % combined
2S
3M
2L
3L
3S
RI45
1C2
ITS1
all
only in H2228
not in H2228
only in H1299-NRF2E79Q
only in H520
only in SKMES1
only in H2228/H520

**Figure S9. NMRAL2P transcript variants across five lung cancer cell lines.** The genomic schematic shows exons 1–6 and the positions of the alternative internal transcription start sites ITS1 and ITS2. Transcript structures detected in A549, H1299, H520, H2228, and SKMES1 cells are shown below. All three major variants were detected in all five cell lines and differ in whether exon 2 is skipped or included with a short (2S) or long (2L) alternative donor site. The bracketed minor variants together account for less than 10% of NMRAL2P transcripts and show additional variation in exon inclusion, splice donor usage, and intron retention. RI45 denotes retention of the intron between exons 4 and 5. Annotations at right indicate the cell lines or conditions in which each variant was detected.

### Figure S10

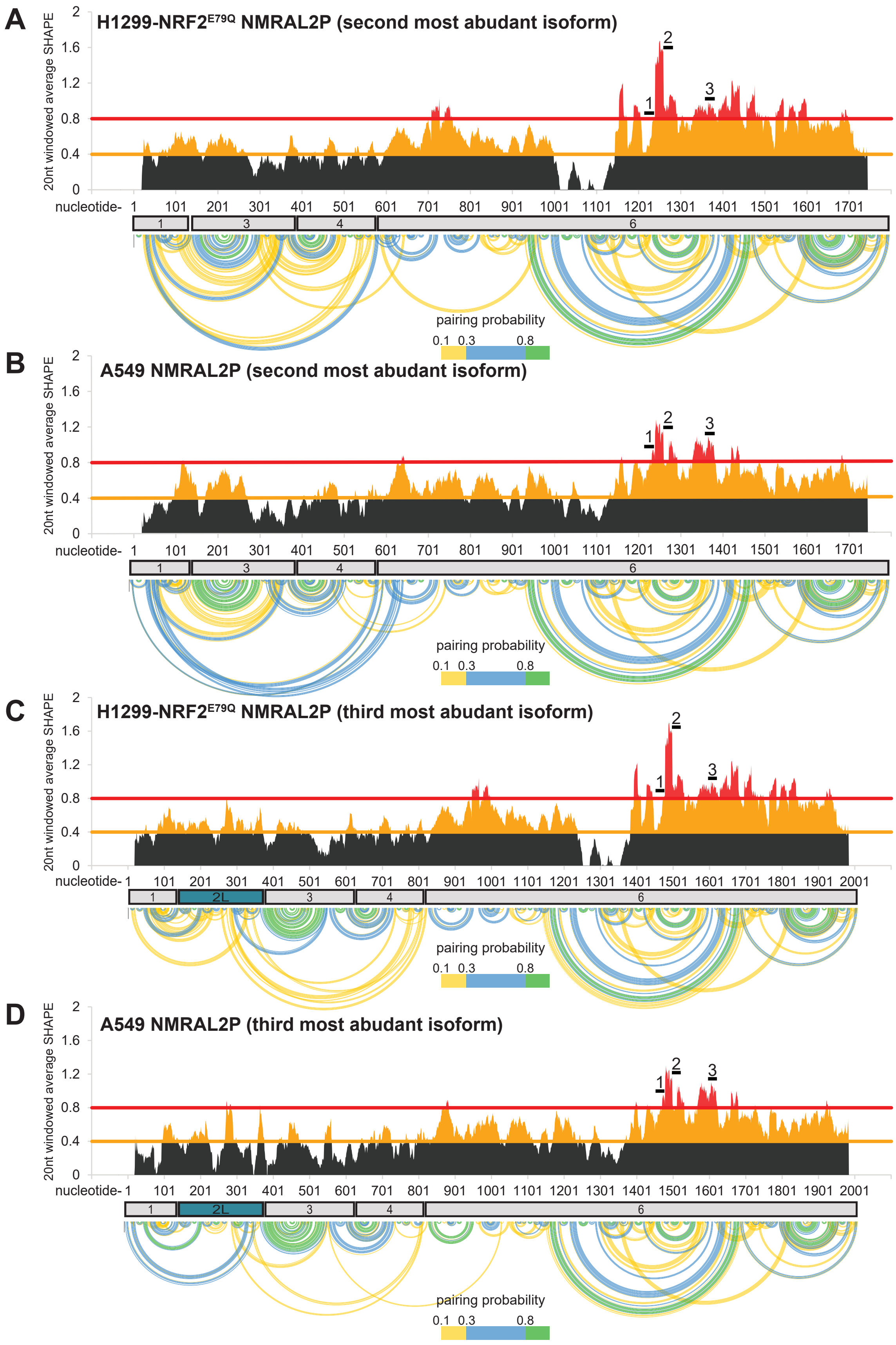
A
H1299-NRF2E79Q NMRAL2P (second most abudant isoform)
20nt windowed average SHAPE
nucleotide-
pairing probability
0.1 0.3 0.8
B
A549 NMRAL2P (second most abudant isoform)
C
H1299-NRF2E79Q NMRAL2P (third most abudant isoform)
D
A549 NMRAL2P (third most abudant isoform)

**Figure S10. SHAPE reactivity and base-pairing probabilities of the second and third most abundant NMRAL2P isoforms.** SHAPE reactivity profiles and predicted base-pairing probabilities are shown for the second most abundant isoform, which skips exon 2, in **(A)** H1299-NRF2$^{E79Q}$ and **(B)** A549 cells, and for the third most abundant isoform, which includes the long form of exon 2 (2L), in **(C)** H1299-NRF2$^{E79Q}$ and **(D)** A549 cells. 20 nt average SHAPE reactivities are plotted by nucleotide position, with low, intermediate, and high reactivities shown in black, orange, and red, respectively. Horizontal lines mark reactivity thresholds of 0.4 and 0.8. Numbered bars above the profiles mark three gapmer targeting sites in exon 6. Boxes below each profile indicate exon boundaries, and arcs connect predicted base pairs, colored by pairing probability as indicated.

#### Figure S11

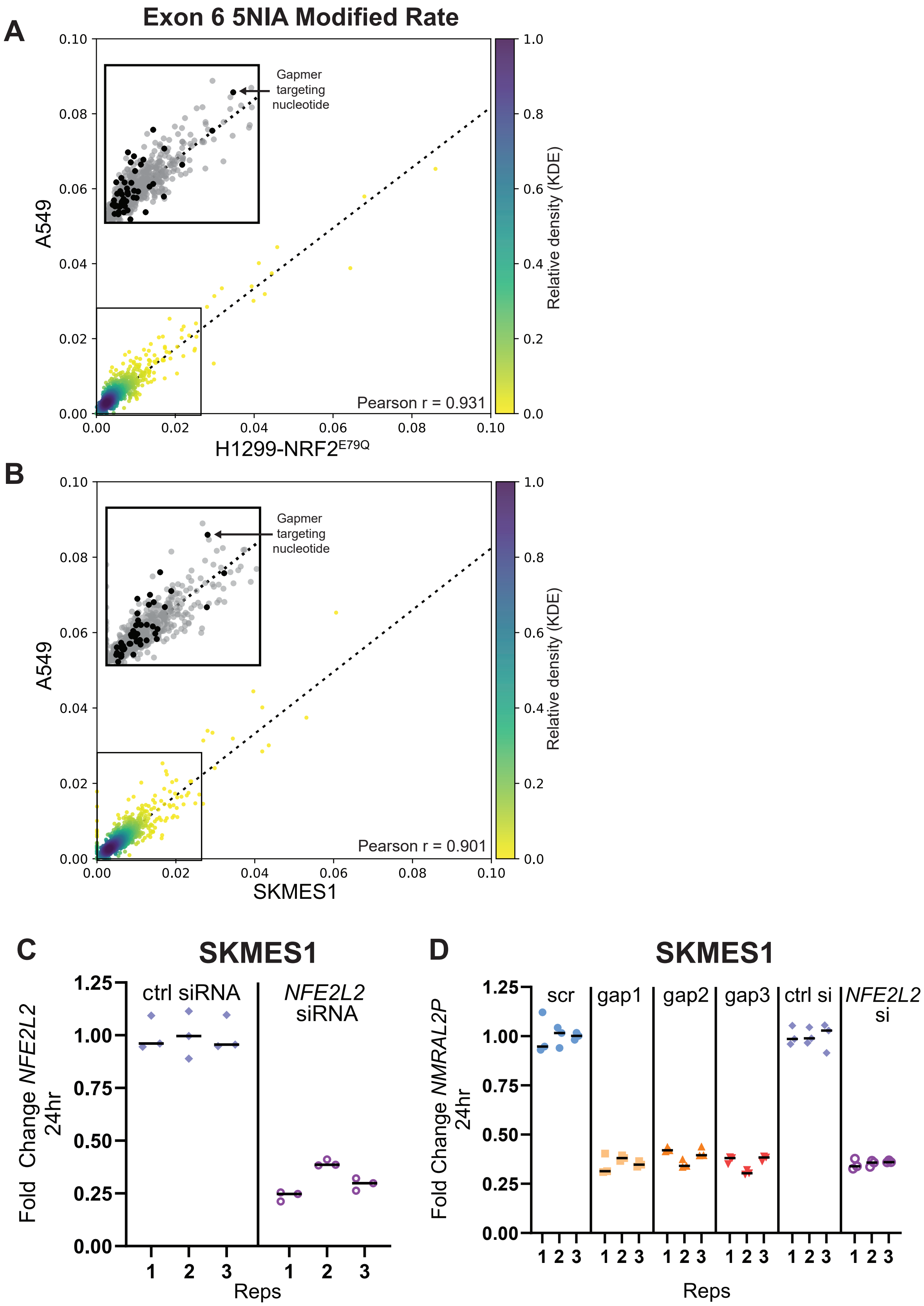
A
Exon 6 5NIA Modified Rate
Gapmer targeting nucleotide
A549
H1299-NRF2E79Q
Pearson r = 0.931
Relative density (KDE)
B
Gapmer targeting nucleotide
A549
SKMES1
Pearson r = 0.901
Relative density (KDE)
C
SKMES1
ctrl siRNA
NFE2L2 siRNA
Fold Change NFE2L2 24hr
Reps
D
SKMES1
scr
gap1
gap2
gap3
ctrl si
NFE2L2 si
Fold Change NMRAL2P 24hr
Reps

**Figure S11. Correlation of NMRAL2P exon 6 SHAPE modification rates and validation of gapmer- and siRNA-mediated knockdown in SKMES1 cells.** Correlation of per-nucleotide 5NIA SHAPE modification rates in NMRAL2P exon 6 between **(A)** A549 and H1299-NRF2$^{E79Q}$ cells and between **(B)** A549 and SKMES1 cells. Insets enlarge the boxed regions and highlight nucleotides within gapmer target sites in black. Point colors in the main plots indicate relative density, and Pearson's *r* is shown for each comparison. **(C)** Quantitative PCR confirmation of *NFE2L2* knockdown in SKMES1 cells 24 hr after transfection with *NFE2L2*-targeting siRNA, compared with control siRNA. **(D)** Confirmation of NMRAL2P depletion in SKMES1 cells 24 hr after transfection with each of three NMRAL2P-targeting gapmers or with *NFE2L2*-targeting siRNA, compared with the scrambled gapmer or control siRNA, respectively. Fold changes are shown for three technical replicates per condition with mean change indicated with black lines.

#### Figure S12

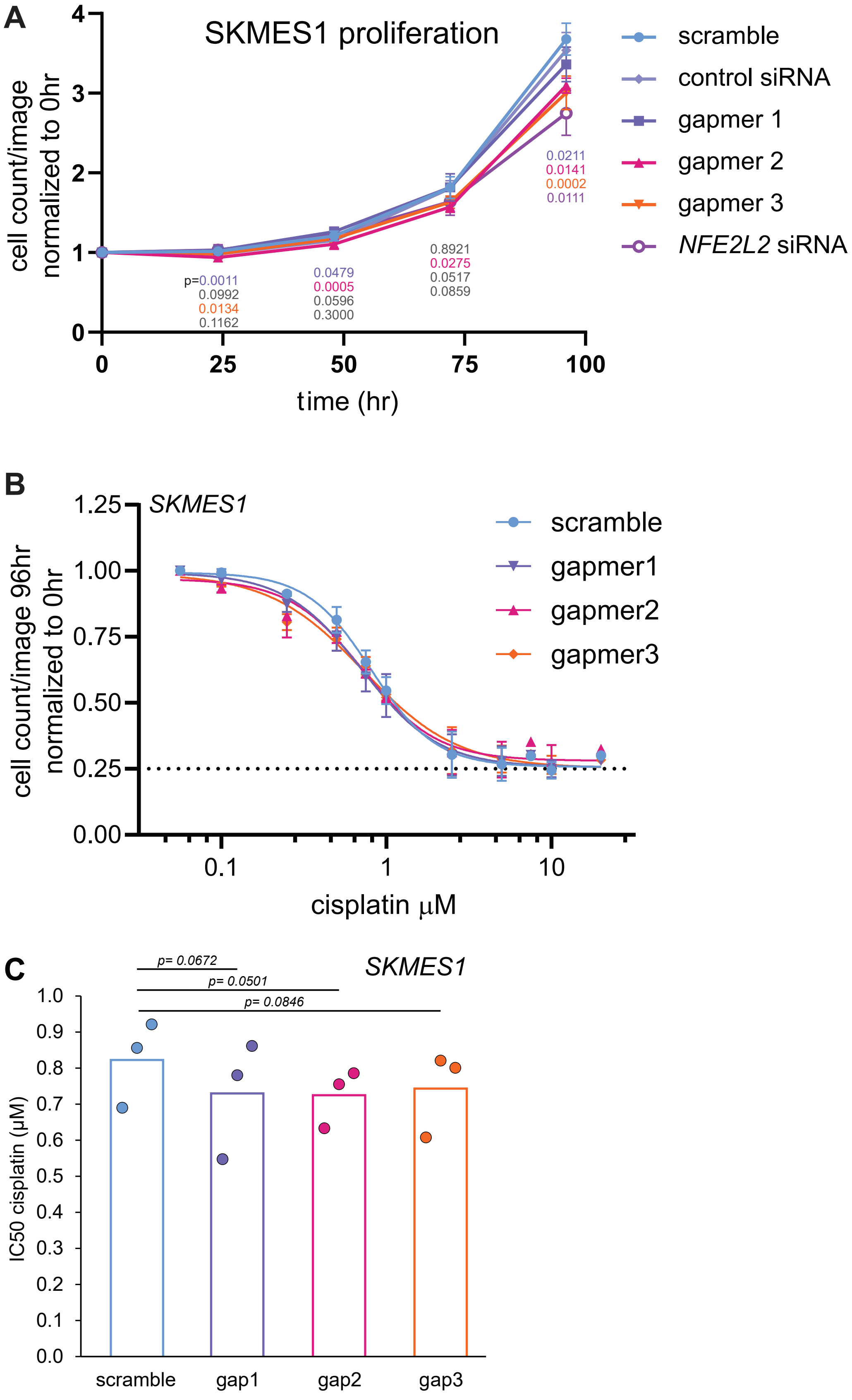
A
SKMES1 proliferation
cell count/image normalized to 0hr
time (hr)
scramble
control siRNA
gapmer 1
gapmer 2
gapmer 3
NFE2L2 siRNA
p=0.0011
0.0992
0.0134
0.1162
0.0479
0.0005
0.0596
0.3000
0.8921
0.0275
0.0517
0.0859
0.0211
0.0141
0.0002
0.0111
B
SKMES1
cell count/image 96hr normalized to 0hr
cisplatin μM
scramble
gapmer1
gapmer2
gapmer3
C
SKMES1
p= 0.0672
p= 0.0501
p= 0.0846
IC50 cisplatin (μM)
scramble
gap1
gap2
gap3

**Figure S12. Effects of NMRAL2P depletion on SKMES1 cell proliferation and cisplatin response. (A)** Proliferation of SKMES1 cells after transfection with one of three NMRAL2P-targeting gapmers, a scrambled gapmer, *NFE2L2*-targeting siRNA, or control siRNA. Cell counts per image were normalized to the corresponding 0 hr counts and tracked through 96 hr. *P* values are shown at the indicated time points. **(B)** Cisplatin concentration-response curves for cells transfected with NMRAL2P-targeting or scrambled gapmers. Cell counts per image at 96 hr were normalized to the corresponding 0 hr counts. **(C)** Cisplatin $IC_{50}$ estimates for each gapmer condition. Points represent three individual replicates, and bars show the mean. *P* values are shown for comparisons with the scrambled gapmer control. All *P* values derive from paired two-tailed t-tests.

### Figure S13

#### RRHO2 pairwise comparison heatmaps

##### A549_gapmers

GAP1 vs Scramble vs GAP2 vs Scramble

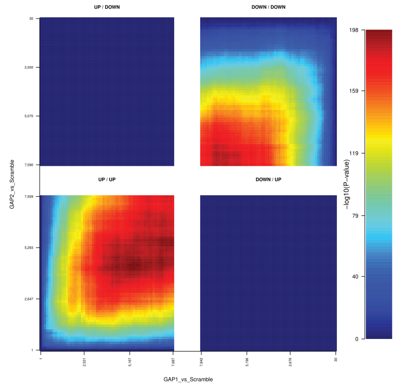


GAP1 vs Scramble vs GAP3 vs Scramble

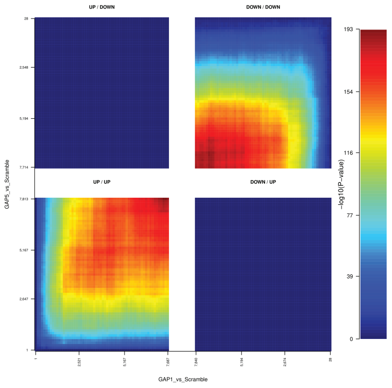


GAP2 vs Scramble vs GAP3 vs Scramble

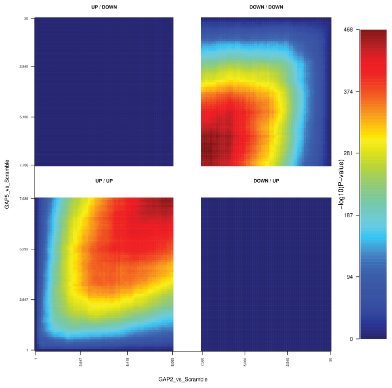


##### SKMES1_gapmers

Gapmer 1 vs Ctrl gapmer vs Gapmer 2 vs Ctrl gapmer

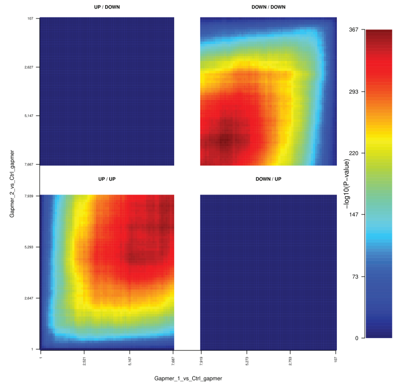


Gapmer 1 vs Ctrl gapmer vs Gapmer 3 vs Ctrl gapmer

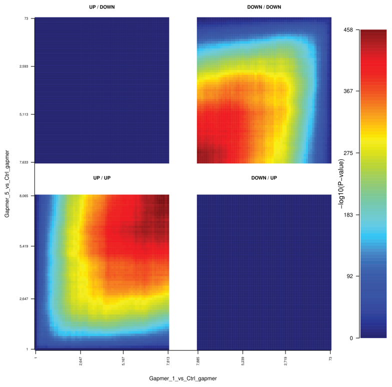


Gapmer 2 vs Ctrl gapmer vs Gapmer 3 vs Ctrl gapmer

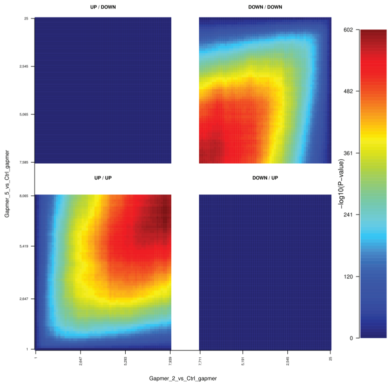


##### NRF2si

NFE2L2 siRNA vs Control siRNA vs NFE2L2 siRNA vs Control siRNA

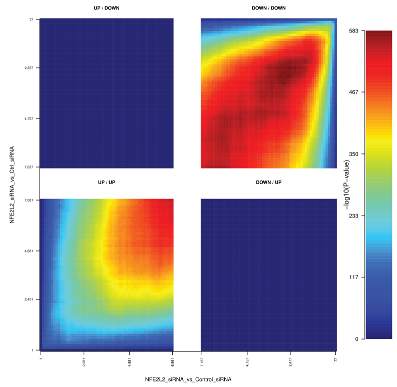

**Figure S13. Pairwise overlap of ranked differential gene expression profiles following NMRAL2P or NRF2 depletion.** Rank-rank hypergeometric overlap (RRHO2) heatmaps compare differential expression profiles across three sets of contrasts. **A549 gapmers.** All three pairwise comparisons among NMRAL2P-targeting gapmers in A549 cells, each profiled relative to the scrambled gapmer control. **SKMES1 gapmers.** All three pairwise comparisons among NMRAL2P-targeting gapmers in SKMES1 cells, each profiled relative to the control gapmer. **NRF2 siRNA.** Comparison of the A549 and SKMES1 responses to *NFE2L2*-targeting siRNA, each profiled relative to its corresponding control siRNA. The axes represent rank cutoffs in the two differential expression profiles. The lower-left and upper-right quadrants show overlap among concordantly upregulated and concordantly downregulated genes, respectively. The upper-left and lower-right quadrants show overlap among genes regulated in opposite directions. Colors indicate $-\log_{10}$ of nominal hypergeometric overlap P values, with a separate color scale for each heatmap.

### Figure S14

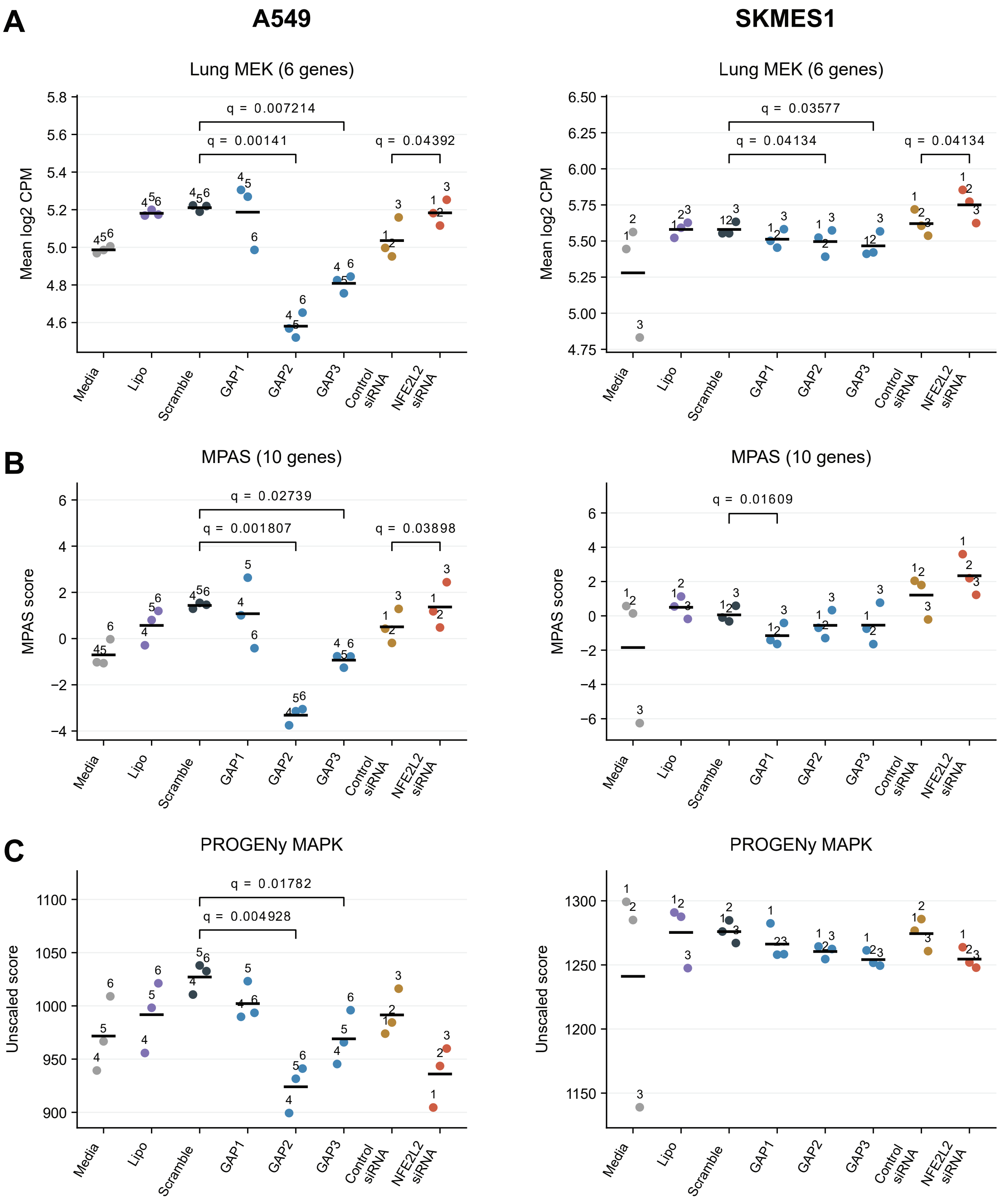

A
A549
SKMES1
Lung MEK (6 genes)
Mean log2 CPM
q = 0.007214
q = 0.00141
q = 0.04392
q = 0.03577
q = 0.04134
q = 0.04134
B
MPAS (10 genes)
MPAS score
q = 0.02739
q = 0.001807
q = 0.03898
q = 0.01609
C
PROGENy MAPK
Unscaled score
q = 0.01782
q = 0.004928
Media
Lipo
Scramble
GAP1
GAP2
GAP3
Control siRNA
NFE2L2 siRNA

**Figure S14. MAPK pathway signature scores following NMRAL2P or NRF2 depletion.** RNA-seq samples from A549 (left) and SKMES1 (right) cells were scored using three MAPK pathway signatures. **(A)** Lung MEK signature score, calculated as the mean $\log_2$ counts per million (CPM) across six genes. **(B)** Ten-gene MAPK pathway activity score (MPAS). **(C)** Unscaled PROGENy MAPK pathway score. Points represent individual replicates, numbers identify replicate blocks, and horizontal bars indicate group means. Each NMRAL2P-targeting gapmer was compared with the scrambled gapmer using a replicate-blocked linear model with shared residual variance across the gapmer groups. *NFE2L2*-targeting siRNA was compared with control siRNA using a two-sided paired t-test. Media and Lipofectamine-only controls are shown for reference. Brackets indicate comparisons with Benjamini-Hochberg-adjusted *P* values ($q$) < 0.05.

### Figure S15

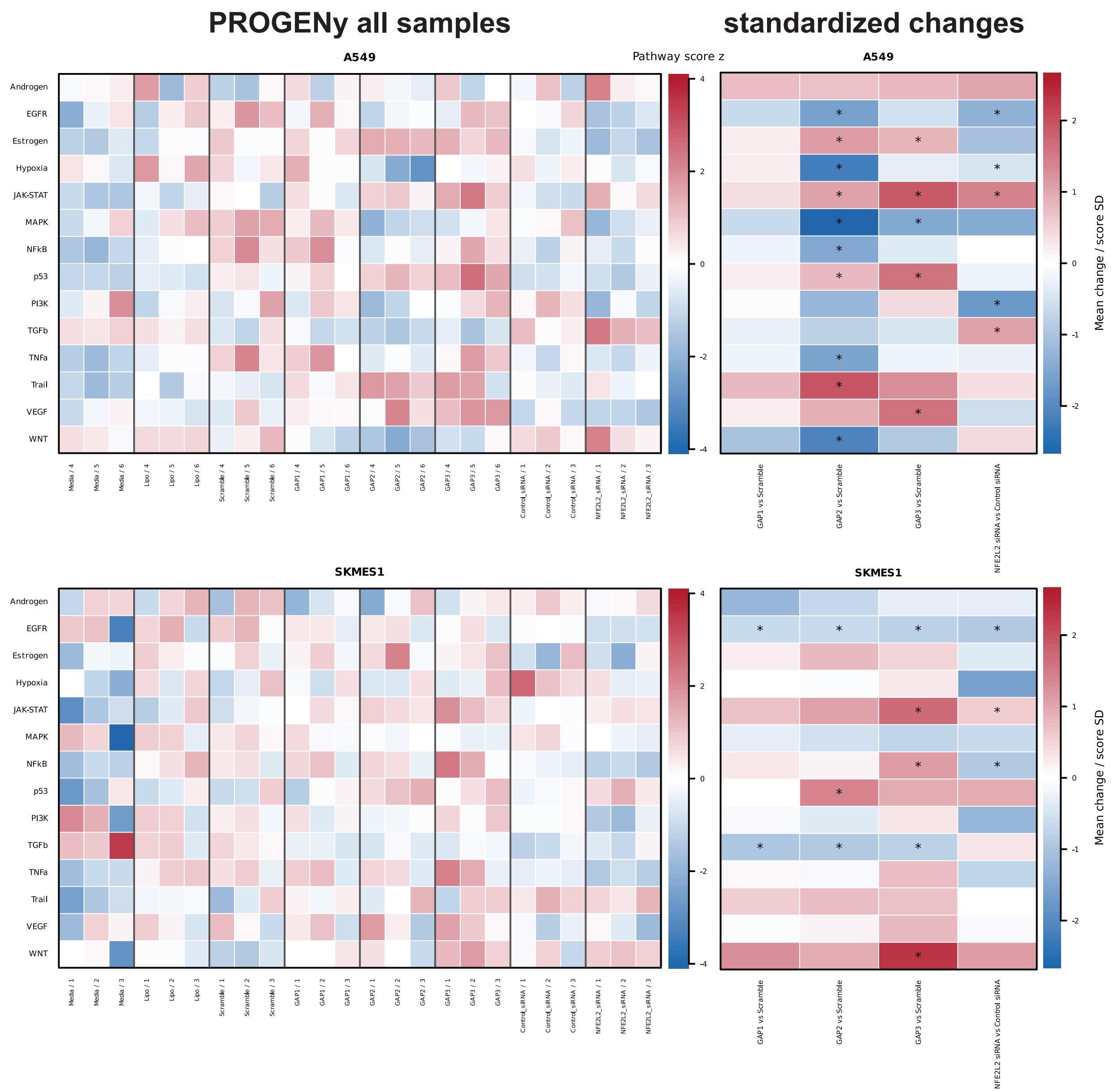

PROGENy all samples
standardized changes
A549
Pathway score z
Mean change / score SD
SKMES1
Androgen
EGFR
Estrogen
Hypoxia
JAK-STAT
MAPK
NFkB
p53
PI3K
TGFb
TNFa
Trail
VEGF
WNT
Media / 4
Media / 5
Media / 6
Lipo / 4
Lipo / 5
Lipo / 6
Scramble / 4
Scramble / 5
Scramble / 6
GAP1 / 4
GAP1 / 5
GAP1 / 6
GAP2 / 4
GAP2 / 5
GAP2 / 6
GAP3 / 4
GAP3 / 5
GAP3 / 6
Control_siRNA / 1
Control_siRNA / 2
Control_siRNA / 3
NFE2L2_siRNA / 1
NFE2L2_siRNA / 2
NFE2L2_siRNA / 3
Media / 1
Media / 2
Media / 3
Lipo / 1
Lipo / 2
Lipo / 3
Scramble / 1
Scramble / 2
Scramble / 3
GAP1 / 1
GAP1 / 2
GAP1 / 3
GAP2 / 1
GAP2 / 2
GAP2 / 3
GAP3 / 1
GAP3 / 2
GAP3 / 3
GAP1 vs Scramble
GAP2 vs Scramble
GAP3 vs Scramble
NFE2L2 siRNA vs Control siRNA

**Figure S15. PROGENy pathway scores following NMRAL2P or NRF2 depletion.** Heatmaps show scores for 14 PROGENy pathways in A549 (top) and SKMES1 (bottom) cells. Left panels show individual sample scores, z-standardized across all 24 samples within each cell line for each pathway. Right panels show the mean treatment-minus-control difference divided by the standard deviation of that pathway's scores across the same 24 samples. Each NMRAL2P-targeting gapmer was compared with the scrambled gapmer using a t-test from a model that matched samples by replicate and pooled residual variance across gapmer conditions. *NFE2L2*-targeting siRNA was compared with control siRNA using a two-sided paired t-test. Red and blue indicate higher and lower scores on the indicated scale. Asterisks indicate Benjamini-Hochberg-adjusted $P < 0.05$, with correction across all 14 pathways and four treatment contrasts within each cell line.
